# Time- and size-resolved bacterial aerosol dynamics in highly polluted air: new clues for haze formation mechanism

**DOI:** 10.1101/513093

**Authors:** Ting Zhang, Xinyue Li, Minfei Wang, Haoxuan Chen, Maosheng Yao

## Abstract

Aerosol chemistry is often studied without considering microbial involvements. Here, we have applied a high-volume (1 m^3^/min) aerosol sampler and the Micro-Orifice Uniform Deposit Impactor (NanoMoudi) along with molecular and microscopic methods to investigate time-and size-resolved bacterial aerosol dynamics in air. Under high particulate matter (PM) polluted episodes, bacterial aerosols were detected to have a viability up to 50-70% in the 0.56-1 μm size range, at which elevated levels of SO4^2−^, NO^3−^ and NH^4+^ were concurrently observed. Engineered or acclimated for both pharmaceuticals and wastewater treatment, bacteria such as *Psychrobacter* spp., *Massilia* spp., *Acinetobacter* lwoffii, *Exiguobacterium* aurantiacum, and *Bacillus* megaterium were shown to have experienced massive abundance shifts in polluted air on early mornings and late afternoons, on which were previously reported to witness rapid new particle formation events. For example, *Acinetobacter* spp. were shown to account for > 96% abundance at a corresponding PM_2.5_ level of 208 ng/m^3^. The bacterial aerosol changes corresponded to the observed PM_2.5_ mass peak shift from 3.2-5.6 μm to the high viability size range of 0.56-1μm. Additionally, it is interesting that elevated levels of soluble Na, Ca, Mg, K, Al, Fe and P elements that are required for bacterial growth were observed to co-occur with those significant bacterial aerosol structure shifts in the air. For particular time-resolved PM_2.5_ pollution episodes, *Acinetobacter* and *Massilia* were shown to alternate in dominating the time-resolved aerosol community structures. The results from a HYSPLIT trajectory model simulation suggested that the role by air mass transport in affecting the observed bacterial aerosol dynamics could be minor. As an evidence, we found that *Acinetobacter, Psychrobacter, Exiguobacterium*, and *Bacillus* genera were emitted into the air with a level of > 3000 CFU/m^3^ from a pharmaceutical plant. In addition, high level of VOCs up to 15,030 ppbv, mainly Acetone (61%) and Acetaldehyde (11%), were also detected in the air inside the plant. All the data including size-resolved viability and time-resolved bacterial aerosol dynamics together with their growth conditions detected in the air suggested that airborne bacteria in the size range of 0.56- 1μm could have played important roles for haze formation in Beijing. The results about time- and size-resolved bacterial aerosol dynamics from this work provide a fresh understanding of aerosol chemistry especially in highly polluted air. It is hoped that these findings could lend a support in future cost-effective air pollution control practices.

## 1. Introduction

Air pollution has become one of serious environmental challenges facing mankind in modern society. Among many others, the core question about air pollution episode is : what is the driving force for accelerating the rapid fine particle growth in typical haze events? Regarding the PM_2.5_ precursor VOC, some studies report that current models underestimate atmospheric VOC emission and also OH radical reactivity(Whalley et al., 2016). Experiments have shown that a direct source of extremely low volatile organic compounds, i.e., a possible new particle formation precursor, was observed from the OH oxidation of acyclic and exocyclic terpenes and isoprene (Jokinen et al., 2015). In another work, it was found that the new particle formation rate, in addition to one-hour lag in time compared to the observation, in the size range of 6-10 nm was significantly underestimated by WRF-Chem and MALTE-BOX models(Huang et al., 2016). These studies collectively imply that there are some uncertainties and unexpected outcomes in measurements of VOCs, new particles and OH reactivity by both aerosol chemistry models and field campaigns.

Although biological materials such as bacteria are increasingly being investigated during haze episodes, their roles in the PM_2.5_ formation have never been studied or considered. Thus, it is completely unclear if the biological materials have a role in the haze occurrence. In the past, during the haze episodes the levels of fluorescent bioaerosol (described to be viable) particles were shown to be elevated in the size ranges of less than 1 μm(Wei et al., 2016). In addition, endotoxin, an important bacterial membrane component, was also shown to be twice that on clear days(Wei et al., 2016). In other studies, culturable bacterial levels were shown to be higher during the haze periods than those during clear days(Liu et al., 2018;Gao et al., 2016;Gao et al., 2015a;Cao et al., 2014). Among the bacterial genera, *Bacillales, Actinomycetales, Pseudomonadales* were detected to dominate the community(Cao et al., 2014;Pöschl et al., 2010). For climate, evidences show that certain bacteria could serve as an ice nucleator or cloud condensation nuclei (Fröhlich-Nowoisky et al., 2016;Šantl-Temkiv et al., 2015;Pöschl et al., 2010), and they might undergo metabolisms even at lower temperatures(Price and Sowers, 2004). Unfortunately, all these studies did not consider the microbial roles in atmospheric aerosol chemistry or the formation of haze episodes.

Here, this work was conducted to investigate the following questions: 1) if certain bacteria experience a rapid massive increase at particular times in highly polluted air?; 2) If so, in which size and what are the species?; 3) Are there any differences in bacterial dynamics during day and night time for a typical haze episode?; 4) If bacterial growth conditions are available in highly polluted air? The answers to these questions will be of great value in providing new insights on the long-sought haze formation mechanism.

## 2. Materials and methods

### 2.1 Particulate matters (PM) collection

#### 2.1.1 PM sample collection using the NanoMoudi

Size-resolved particulate matter samples were collected using Micro-Orifice Uniform Deposit Impactor (NanoMoudi) (MSP, USA) with 13 stages (stage 1: 10-18 μm; stage 2: 5.6-10 μm;stage 3: 3.2-5.6 μm; stage 4: 1.8-3.2 μm; stage 5: 1.0-1.8 μm; stage 6: 0.56-1.0 μm; stage 7: 0.32-0.56 μm; stage 8: 0.18-0.32 μm; stage 9: 0.1-0.18 μm; stage 10: 0.056-0.01 μm; stage 11: 0.032-0.056 μm; stage 12: 0.018-0.032 μm; and stage 13: 0.01-0.018 μm) near the Environmental Building on Peking University campus(40°00′N, 116°32′E). For each experiment, the NanoMoudi at a sampling flow rate of 28 L/min was placed at 1.5 m above the ground. For the NanoMoudi experiment, six sets of samples were obtained on 47 mm (stage 1 to 10) and 90 mm (stage 11 to 13) aluminum membranes from Sep. 11, 2017 to Sep.14, 2017, during which a high PM_2.5_ pollution episode occurred. The sampling time was categorized into the day-time (Day): from 7:00 AM to 7:00 PM, and the night-time (Night): from 8:00 PM to 6:00 AM.

Before and after sampling, aluminum membranes on each stage of the sampling device were weighed using a microbalance after conditioned at constant temperature (24.5°C) and humid (50%) for at least 48 h in advance. The membrane with samples on each stage of the NanoMoudi was then placed into a 15 mL tube and soaked in 4 mL 0.05% tween 20 water, and the sealed tubes were placed at 4°C in a shaker operated with a rotation speed of 200 rpm continuously for 4 hours. Finally, all the well-mixed sample extracts were kept at −20°C immediately before any analysis. For the quality controls, all aluminum membranes and the sampler were washed by absolute ethyl alcohol before use. And blank membranes without the air sampling were also brought to the sampling site which was later incorporated into the sample analysis as the negative control.

### 2.2 Time-resolved bacterial aerosol samples collected using the HighBioTrap sampler

Between February 2018 and May 2018, air samples were collected for a 24-hr time period for each day in three different pollution episodes (14 −208 ng/m^3^) using the HighBioTrap sampler (Beijing dBlue Tech, Inc, Beijing, China) at an air flow rate of 1000 L/min for 20 min for each sample on Peking University campus (40°00′N, 116°32′E). The detailed sampling protocol was described for the HighBioTrap in our previous work(Chen and Yao, 2018). For each PM_2.5_ pollution episode, particulate matter samples were collected in duplicates using the HighBioTrap at an interval of every 2 or 4 hours on an aluminum membrane coated with 600 μL mineral oil at a height of 1.5 m above the ground. Specific sampling times and meteorological conditions are provided in the Table S1 (Supporting Information). Here, two air samples with 20 m^3^ air were collected using the high-volume sampler for each specific time period. As a comparison, bacterial aerosol samples were also collected using the HighBioTrap in a pharmaceutical plant to study the bacterial emissions into the air. After each sampling, the mineral-oil-coated membrane was washed by 1.5 mL 0.05% tween 20 water, then the oil-in-water emulsion was centrifuged to remove the mineral oil supernatant at a speed of 7000 rpm using a centrifuge (5804r, Eppendorf, German). After a mixing process, the sample extracts were kept at −20°C immediately before any analysis. For total bacteria and their community structures, 78 samples and 2 negative control samples (blank membrane samples) were analyzed using qPCR and Illumina platforms (Sangon Biotech, Inc., Shanghai, China) as described in in Supplementary Text. Besides, bacterial species identifications were also performed using VITKE MS and MICROFLEX as described in Supplementary Text. As a comparison, air samples were also collected using the 4-Channel Particulate Matter Sampler as documented in Supplementary Text.

Besides, air samples from a pharmaceutical plant were also collected using Silonite^™^ Classical Canisters (Entech Instruments, Simi Valley, CA 93065), and their corresponding volatile organic compounds (VOCs) levels were further analyzed (Wuhan Tianhong Instruments Co., Ltd.) according to a previously published protocol^34^.

### 2.3 PM-borne *Endotoxin assay using the LAL assay, bacterial viability using DNA stain method, and reactive oxygen species (ROS) using DL-Dithiothreitol (DTT) assay*

The samples collected from the NanoMoudi were first centrifuged at 4°C at 3000 rpm in 10 min, and the supernatants were then transferred to the new tubes for the Limulus amebocyte lysate (LAL) assay. A 5-point standard curve from 0.005 to 50.0 EU/ml was generated using 10-fold serial dilutions of endotoxin standards (Control Standard Endotoxin, CSE, Associate of Cape Cod, Inc., Eastham, MA, USA) by endotoxin-free water (LAL Reagent Water, Associate of Cape Cod, Inc., Eastham, MA, USA). The LAL agent (Associate of Cape Cod, Inc., Eastham, MA, USA) diluted in the Glucashield Beta Glucan Inhibiting Buffer (Associate of Cape Cod, Inc., Eastham, MA, USA) was used in the assay according to the manufacturer’s guidelines. The standards, negative controls and air samples were pre-incubated and analyzed using a microplate reader running SoftMaxPro 5.4.1 software (SpectraMax 340; Molecular Devices, Sunnyvale, CA) with photometric measurements taken at 37°C every 1min for 60 min at 405 nm. The viability of bacteria in the particulate matters was studied using a live/dead viability kit (L7012 BacLight Viability Kit, Invitrogen) as described in Supplementary Text. PM-borne reactive oxygen species (ROS) and metal analysis for certain air samples collected were conducted using DTT assay and ICP-MS as documented in Supplementary Text.

### 2.4 Air mass back-trajectory analysis with cluster

Air mass transport for three-days was studied during certain air pollution episodes using HYSPLIT Trajectory Model(NOAA) via Hysplit 4 software. The climate data from GDAS1(ftp://arlftp.arlhq.noaa.gov/pub/archives/gdas1/) were used for simulation.

## 3. Statistical analysis

The mass distribution mode differences between the hazy day and clean day were analyzed by two-way ANOVA. Other difference analyses were performed using the paired t-test. All tests were performed using the statistical components from the SigmaPlot 12.5 software. Non-metric Multi-Dimensional Scaling (NMDS) based on Bray-Curtis distance was used to compare bacterial community structures among samples. NMDS was performed by *R* software (version 3.2) with the vegan package 2.0-10. A p-value of 0.05 indicates a statistically significant difference in this work.

## 4. Results and discussion

### 4.1 Size-resolved PM mass and composition during different pollution episodes

Here, we presented our results about PM loading and bacterial aerosol increases in the air from our 24-hr monitoring of four high PM pollution episodes in Beijing during 2017-2018. On Sept 11-14, 2017, in contrast to peak levels in the size range of 3.2 to 5.6 μm on clear days (PM_2.5_=21.42 μg/m^3^), significant PM_2.5_ fraction increases from less than 30% to up to 90% of the total mass were detected in the size range of 0.56 to 1 |am during the haze episodes (PM_2.5_=91.33 ng/m^3^) as observed in Figure 1 (A), seconded by those appearing in 0.32-0.56 μm size range. For particles of 0.24-0.74 μm, their mass percentages all increased (p-value<0.001; two-way ANOVA), but others did not decrease significantly (p-value=0.198-0.64) as seen in Figure 1 (A) during the haze episodes. The data observed in Figure 1 (A) are generally in line with other studies in which the same particle size range, i.e., 0.56-1 μm, was detected to have the highest mass increases during typical high PM pollution episodes in Beijing (Sun et al., 2015;Yue et al., 2018). This phenomenon was also observed during the haze episodes in other places, such as Taiwan(Tsai et al., 2012) and Singapore(Behera et al., 2015), though different from the firework-caused air pollution scenario(Jing et al., 2014). Similarly, we have also observed that for both size ranges (0.32-0.56 μm and 0.56-1 μm) metals such as Al, Fe, Mg and Zn have all increased during the haze days (B-day and C-night) (Supporting Information Figure S1). For metal Al, Fe and Zn, more emissions were found in the size range of 0.56 - 1 μm during the day than during the night as shown in Figure S1. Particularly, during the haze episodes (B-day and C-night) high Fe levels were detected in 0.32-0.56 μm size range during the night as observed in Figure S1.

**Figure 1.**
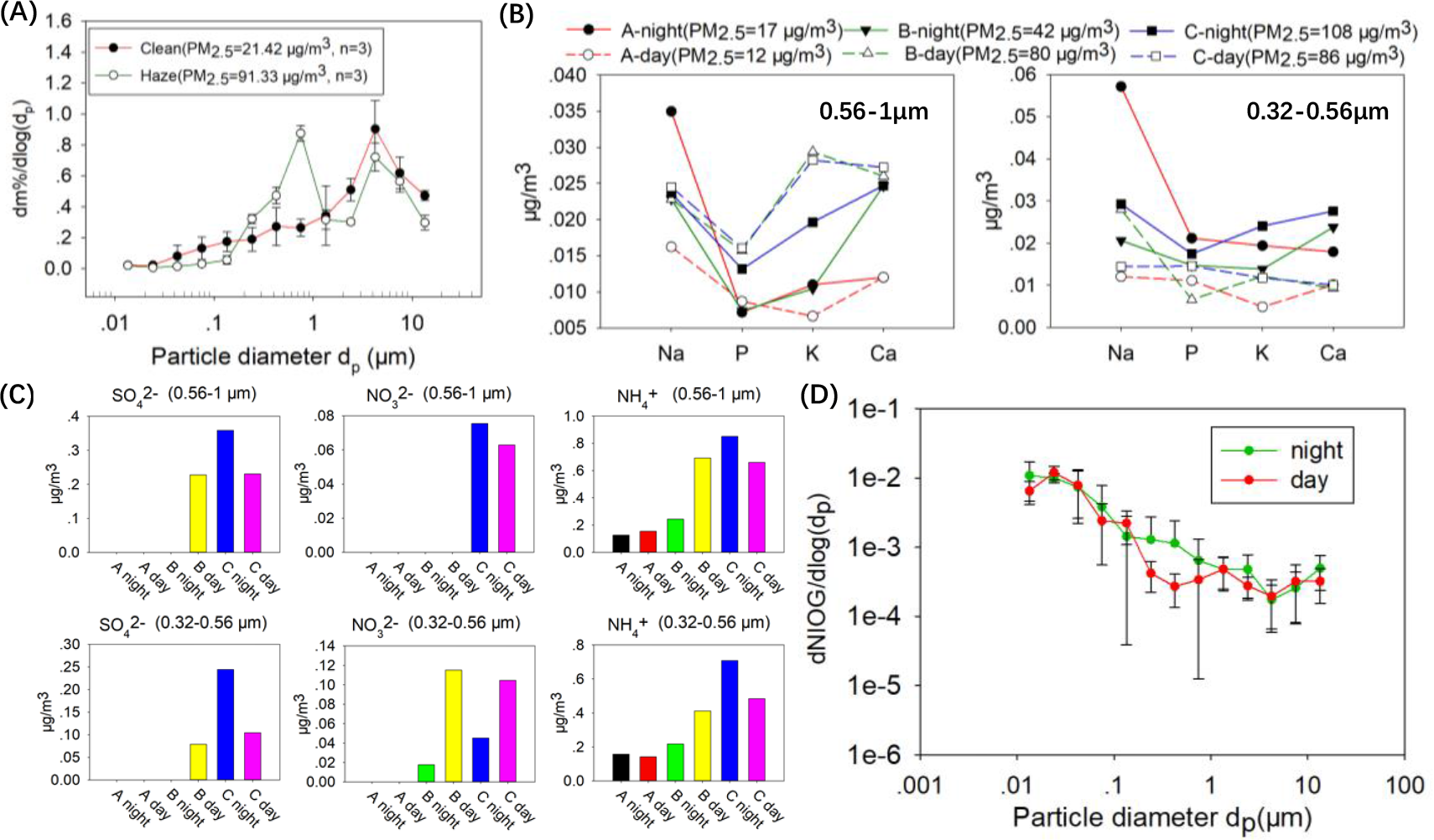
Rapid particle increase during typical high PM pollution episodes on Sept 1114, 2017: A) size-resolved particle mass percentage distribution during low (21.42 pg/m^3^) and high (91.33 pg/m^3^) pollution levels (PM_2.5_ values are averages of three different days (Sept 11-14, 2017); B) Na, P, K, Ca concentration levels in particles of 0.56-1 pm and 0.32-0.56 pm during different time periods (Day: 7:00AM-19:00PM; Night: 20:00PM-6:00AM on Sept 11-14, 2017) under different PM pollution levels (PM_2.5_=12-108 pg/m^3^); C) SO_4_^2−^, NO_3_^−^, and NH_4_^+^ concentration levels measured using Ion Chromatography (Thermo Scientific, Dionex, ICS 2000 and ICS 2500) in particles of 0.56-1 pm and 0.32-0.56 pm under different PM pollution levels on Sept 11-14, 2017; (Corresponding O_3_, NO_2_, SO_2_, CO concentration levels are shown in Supporting Information Figure S2); D) Size-resolved (10 nm-20 μm) particle toxicity measured using DTT method during night and day as shown in Figure 1 (B). Air samples were collected using the NanoMoudi sampler at a flow rate of 28 L/min for 12 hours during the Day (7:00 AM to 19:00PM) and 10 hours during the Night (20:00PM to 6:00AM) on Sept 11-13, 2017.

For other trace metals as shown in Supporting Information (Figure S3), two size ranges have not experienced major changes. As for metal As, significant elevation was detected during the daytime, especially in the size range of 0.32 to 0.56 μm during the haze episode. For the C-Day as referred in Figure 1, the metal As (coal combustion indicator) seemed to have shifted from size of 0.32-0.56 μm during the night to the size of 0.56-1 μm during the day. For Ni, even for the clear day (A) higher concentration levels were detected in the size range 0.56-1 μm, and no elevated levels were detected during the haze episodes, suggesting that Ni contribution from fugitive dusts or industrial activities might be insignificant for the haze formation studied in this work. For both size ranges (0.32-0.56 μm) and (0.56-1 μm), metals such as Mn, Cu, Se, Ba and Pb were detected to have higher levels than those during the clear day. As can be seen from the figure, during the nighttime higher element Se levels were detected for the C-day (higher PM_2.5_ levels) in the size range of 0.32 to 0.56 μm, however during the daytime peak Se levels were found in the size range of 0.56-1 μm, suggesting Se-containing particles have possibly shifted from smaller sizes to larger sizes. Se is an indicator for coal combustion, and its elevated levels in the particle peak size suggest the coal combustion was one of the driving factors for the haze formation.

To further study PM_2.5_ formation during polluted days, we analyzed the K, P, and Ca elements from bio-mass burning in the PM samples for different size ranges as shown in Figure 1 (B) and Figure S3. The data in these figures suggest that the elements such as K might have shifted to different size ranges during the dynamic aerosol processes. Here, we have also studied SO_4_^3-^, NO_3_^−^ and NH_4_^+^ concentration levels in the particles of 0.32-0.56 μm and 0.56-1 μm, and as observed in Figure 1 (C) their concentration levels were all elevated during the high pollution episodes (e.g., the C-day with higher NO_2_ levels as shown in Figure S2 (Supporting Information)) in contrast to low levels (e.g., the A-day). Literature data show that during a typical haze episode of Beijing sulfate production (Figure S2 showing a higher SO_2_ level between 13 and 14, Sept, 2017) is a major contributor to PM_2.5_ increase(Wang et al., 2016). Here, we also investigated the toxicity of size-resolved PM samples collected from different time periods (daytime and nighttime) and found as shown in Figure 1 (D) that for the size ranges of 0.24-0.74 μm the average PM_2.5_ toxicity (oxidative potential) was higher during the night than during the day though at a lower confidence level (p-value=0. 190), suggesting possible particle composition change during the day. For example, metals such as Fe and As, contributors to PM oxidative potential, in smaller sizes were observed here to have shifted to larger ones. Overall, the observed particle loadings and related chemical compositions here agree with those reported in the literature.

### 4.2 Massive changes in time-resolved bacterial aerosol concentration and viability in highly polluted air

Importantly, we have here observed pronounced bacterial aerosol changes in viability for highly polluted days C-night (PM_2.5_=108 μg/m^3^) and B-day (PM_2.5_=80 μg/m^3^) as shown using viable bacterial percentages in Figure 2 (A) and Supporting Information (Figure S4, S5). Here, we have simultaneously observed the elevated levels of bacterial growth materials such as ions (NO_3_^−^, NH_4_, etc) as shown in Figure 1 (C) at the same size ranges as discussed above, which co-occurred with increases in bacterial cells and viability as shown in Figure 2 (A). Through fluorescent microscopic analysis of the size-resolved PM samples collected using the NanoMoudi as shown in Figure S4, S5 (Supporting Information), we have found that the size-resolved bacterial percentages have varied over the time. For example, as shown in Figure 2 (A) during the low PM_2.5_ day (A-day) most of the bacteria were in the larger size ranges (larger than 1 μm) during the nighttime, however during the daytime the peak (about 22%) shifted to the size range of 0.56-1 μm. Likewise, similar findings were observed for the high PM_2.5_ day (B) except the percentage was detected to be up to 35%. In contrast, for the high PM_2.5_ day (C) the peak significantly decreased from 0.56-1 μm to larger sizes from the nighttime to daytime. The C-day witnessed a decrease of PM_2.5_ levels. For both A-Day and C-Day, the total bacteria have decreased from nighttime to daytime, however for the B-day the total bacteria increased from nighttime to daytime. In addition to microscopic analysis, we have also performed qPCR analysis for the size-resolved bacterial aerosol samples as shown in Figure S6 (Supporting Information). Overall, the qPCR data agreed well with the microscopic data (Supporting Information Figure S4). The total bacteria seemed to have decreased from nighttime to daytime, however for a particular size range, e.g., 0.56-1 μm, their percentages have increased from nighttime to daytime. Overall, these data revealed that bacterial levels in the size ranges of 0.56-1 μm have changed from nighttime to daytime, implying a changing bacterial aerosol dynamic under different time and pollution conditions.

**Figure 2.**
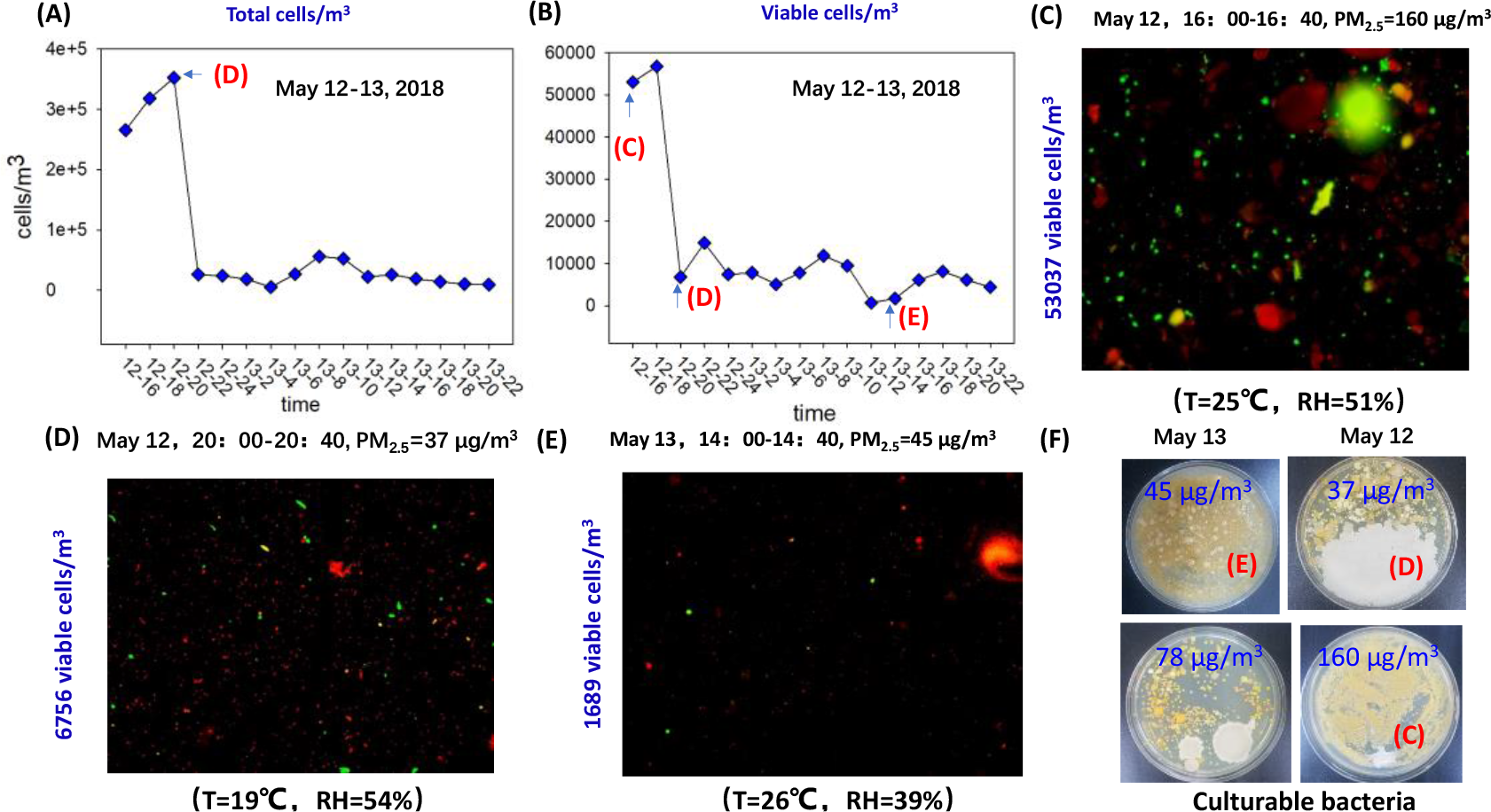
Time-resolved bacterial aerosol dynamics in highly polluted air: A) Total bacterial aerosol concentration levels detected at different times on May 12-13, 2018 using qPCR; B) Total viable bacterial aerosol particle concentration measured using BackLight DNA stain method for different time periods (the digit separated by “-” represent date and time, respectively, e.g., “12-22” means 22:00PM on May 12) during May 12-13, 2018 under different air pollution levels (37 −160 ng/m^3^) (PM_2.5_ levels for different time periods of May 12-13 are presented in Supporting Information Figure S8); C) Example image of bacterial DNA stain results from air samples collected during a high PM_2.5_ pollution episode (PM_2.5_=160 ng/m^3^ at 20:00-20:40 on May 12, 2018); D) Example image of bacterial DNA stain results from air samples collected during a low level PM_2.5_ pollution episode (PM_2.5_=37 μg/m^3^ at 16:00-16:40 on May 12, 2018); E) Bacterial DNA stain results from air samples collected during a medium level PM_2.5_ pollution episode (PM_2.5_=45 μg/m^3^ at 14:00-14:40 on May 13, 2018); F) Example images of bacterial aerosol culturing agar plates (1.33 m^3^ air collected using the HighBioTrap) under different PM_2.5_ mass levels during May 12-13, 2018.

Using BackLight DNA stain method, 50-70% of viable bacteria were found in the size range of 0.56-1 μm as observed in Figure 2 (B) and Figure S4, S5 for the A-day and B-day (haze formation day) during the daytime, while less than 20% of viable bacteria for the C-day (haze disappearing day) was noted. In contrast, during the nighttime, the peak for viable bacteria about 30% was found at 0.32-0.56 μm for B-night, close to 60% found at the 0.56-1 μm for C-night and about 10% for the A-night (Supporting Information Figure S5). The size-resolved percentages of viable bacteria for three different days (A-“biomass burning” day, B- “haze formation”day, C-“haze disappearing” day) were observed to have changed substantially (Supporting Information Figure S5). For dead bacteria, as shown in the figure, in the size range of 0.56-1 μm, higher percentage was observed for the C-day, and the lowest for the B-day (the haze formation day) during the nighttime. However, for the daytime, the highest percentage was observed in the size range of 5.6-10 μm for the C-day (haze disappearing day). The percentages for the A- and B-day were found to be similar as seen in the figure. In this work, we have also analyzed the endotoxin (a bacterial derivative) levels during the nighttime and daytime for three different dates (A, B, and C) as shown in Supporting Information Figure S7. As shown in the figure, for the A- and B-day most of the endotoxin was detected in the smaller size ranges of 0.032 −0.18 μm, while during the daytime most of the endotoxin was detected in the larger sizes of > 3.2 μm. In this work, we have also observed that the endotoxin levels for those hazy days (B and C) were found to be higher than the clear day (A) with lower PM_2.5_ levels for 1-18 μm size range (p-value=0.023), although no statistically significant differences were detected between daytime and nighttime. In a previous work, Wei et al. also reported that endotoxin concentration in the PM was elevated up to 12.4 EU/m^3^ during a haze episode in Beijing, about twice that during a clear day(Wei et al., 2016). All the data including the microscopic data (size-resolved percentages of viable bacteria, qPCR, viability and endotoxin, total bacteria) together with favorable bacterial growth conditions suggest that bacteria in the size range of 0.56− 1μm could have played a non-neglected role for air pollution episodes.

To further understand the problem with bacteria in highly polluted air, we have conducted another three different sets of experiments on May 12-13, 2018, March 10-12, 2018, and Feb 6-7, 2018. Different from the first NanoMoudi experiment, we took time-resolved air samples (20 m^3^ air for 20 min) using a high-volume sampler (HighBioTrap, dBlueTech Co., Ltd, Beijing China) every two hours for different time periods to capture the bacterial growth events. As seen from Figure 2 (A, B, C), at 16:00 of May 12 with a PM_2.5_ level up to 160 ng/m^3^, we have detected a significantly higher bacterial aerosol concentration using qPCR, which also corresponded to higher viable bacterial levels (53,037 cells/m^3^) as detected using DNA stain (Figure 2 (C)). For other time periods, the bacterial aerosol concentration levels were detected to decrease substantially (2 hours after the high PM episode) as shown in Figure 2 (B) down to 1689 cells/m^3^ as shown in Figure 2 (A) by qPCR; Figure 2 (B) and Figure 2 (E) by DNA stain, respectively for different PM_2.5_ levels. Figure 2 (F) shows the culturable bacteria for 1.33 m^3^ air sampled during different PM_2.5_ levels, and the results were comparable to those detected using DNA stain method as shown in Figure 2 (C), Figure 2 (D) and Figure 2 (E). As observed in Figure 2 (F), substantially higher bacterial concentration levels were observed for higher PM_2.5_ levels (160 μg/m^3^). Wei et al. reported that the viable bioaerosol particle concentration increased over a 6-fold at night or early dawn during the haze episode compared with that on clean day(Wei et al., 2016). Culturable bacteria concentrations during the haze days were also observed to be higher than those during the non-haze days(Li et al., 2015). In another work aerosolized bacteria were shown to experience metabolic activity in the simulated air state when supplied with carbon source (Krumins et al., 2014). However, Amato et al. reported that after the 18h aerosolization of the bacteria into the air its cultivable bacterial percentage if without adding nutrients started to decrease down to 4% (Amato et al., 2015). Our data from qPCR, DNA stain, and culturing in these sets of experiments as illustrated in Figure 2 further reveal that massive changes in time-resolved bacterial aerosol dynamics are taking place in highly polluted air, e.g., during haze episodes in Beijing.

### 4.3 Massive time-resolved bacterial community structure shifts in highly polluted air

To further investigate the specific bacterial species that have experienced the massive changes in the polluted air, the collected air samples were sequenced and also analyzed using ion chromatography methods (both VITEK MS and Microflex). Here, we have found as shown in Figure 3 (A) that during clear days (14 to 93 μg/m^3^) as a control on Feb 6-7, 2018 regardless of the sampling time periods, the bacterial structures remained to be relatively uniform for most of the studied time. Nonetheless, we have still detected a difference in bacterial structures as seen in Figure 3 (A) for the time of 20:00PM on Feb 6, 2018, showing an increase in abundances of *Ralstonia* and *Acinetobacter*. Interestingly, this event also co-occurred with a higher PM_2.5_ level, i.e., 75 ng/m^3^. For the clear day, among those bacterial genera detected, *Ralstonia* and *Acinetobacter* were found to be most abundant. Besides, *Psychrobacter* and *Massilia* were also detected among the top 20 genera. In contrast, as shown in Figure 3 (B) the bacterial structures varied greatly within a day during the haze days (PM_2.5_ level up to 251 ng/m^3^) on March 10-12, 2018. Among the bacterial genera detected, *Psychrobacter, Massilia, Acinetobacter*, and *Arthrobacter* were found to dominate the community structure. During low level of air pollution episodes as shown in Figure 3 (A), *Ralstonia* was detected to be the second dominant species up to 4.16-14.66%, however during high pollution episodes (PM_2.5_ concentration up to 251 μg/m^3^), *Ralstonia* was found to only account for 0-0.78% of the total abundances, a substantial decrease. For certain time periods (from 8:00 AM on March 11 to 12:00 AM on March 13), as observed in Figure 3 (B) when the PM_2.5_ levels increased the relative abundances of *Psychrobacter, Massilia* and *Acinetobacter* witnessed sharp increases up to about 96% at the corresponding PM_2.5_ level of 208 ng/m^3^. The bacterial community structure differences are much visible as seen by the genus heatmap shown in Figure 3 (C) for March 10-12, 2018.

**Figure 3.**
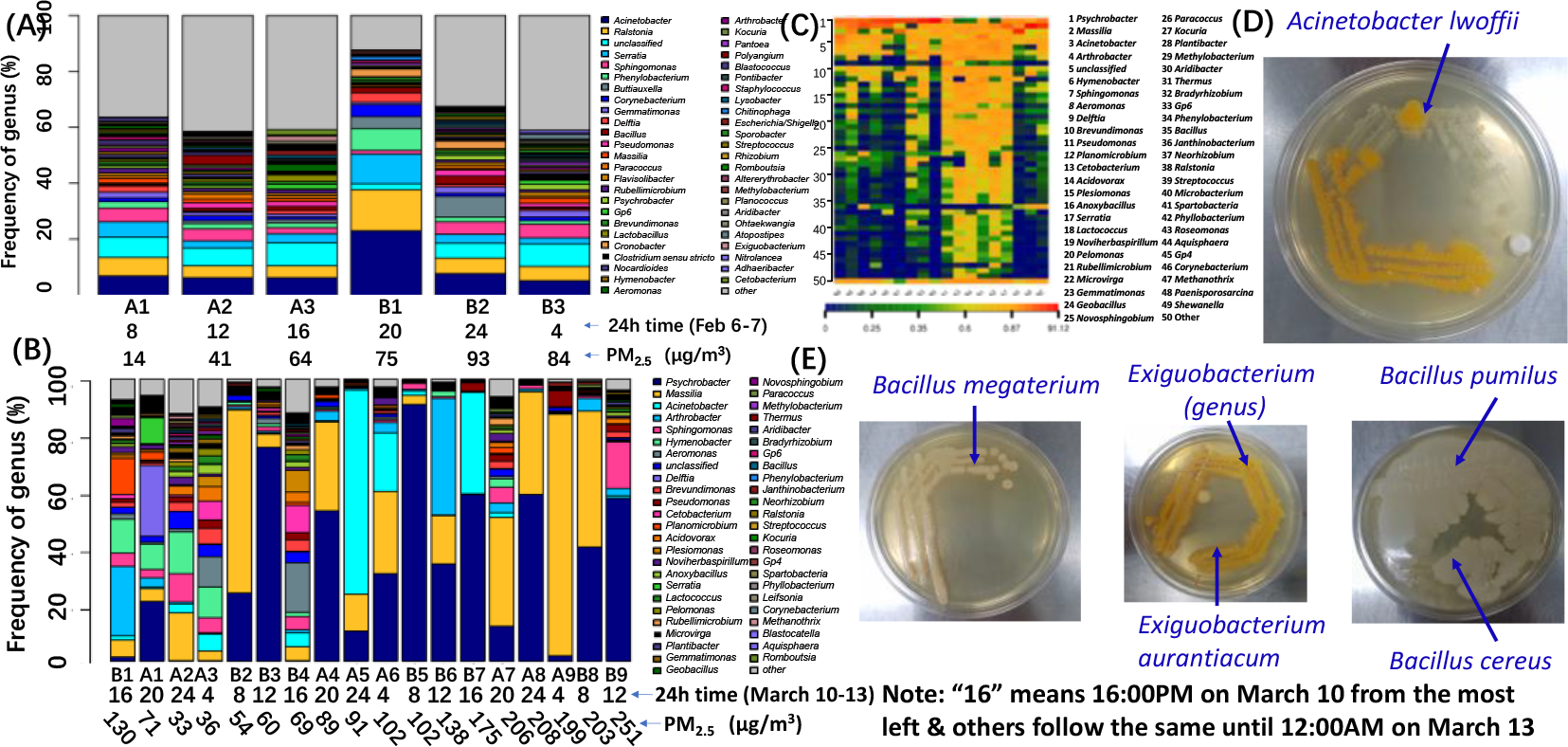
Dynamics of bacterial community structures for different time periods during typical pollution processes: (A) Bacterial community structures (on Feb 6-7, 2018) with PM_2.5_ levels from 14 to 93 ng/m^3^; (B) Bacterial community structures (from March 10 to 12, 2018) with PM_2.5_ levels from 33 to 251 ng/m^3^; (C) Heatmap of bacterial genera structures as shown in Figure 3 (B) on March 10-12, 2018; (D) Culturable bacterial isolates detected during May 12, 2018 high PM_2.5_ pollution episodes as shown in Figure 2; Two air samples were collected using the HighBioTrap for 20 min each at a flow rate of 1000 L/min, and the samples after the culturing were analyzed using VITKE MS (BioMérieux) and microflex (Bruker) for their corresponding bacterial species; *Acinetobacter Iwoffii* and *Exiguobacterium aurantiacum* were found to be most abundant.

To further investigate the problem and confirm these findings, we have conducted another set of different experiments on May 12, 2018 using similar methods. In contrast to other experiments, culturable *Acinetobacter lwoffii, Bacillus cereus, Bacillus pumilus*, *Exiguobacterium aurantiacum*, *Arthrobacter ilicis*, *Bacillus cereus*, *Bacillus megaterium, Coriobacterium* were detected to be abundant using colorimetry methods including VITKE MS (BioMérieux) and microflex (Bruker) (two methods confirmed with each other for specific species) as seen in Figure 3 (D) and Figure 3 (E). Among these species, *Acinetobacter lwoffii, Exiguobacterium aurantiacum,* and *Bacillus pumilus* were found to be most abundant. *Bacillus pumilus* and *Exiguobacterium aurantiacum* are often used in wastewater treatment practices, while *Acinetobacter lwoffii* is a human pathogen. The differences observed in bacterial aerosol dynamics in different PM pollution episodes indicate that air pollution episodes could varied from one to another in their characteristics. For low and high PM pollution levels, the bacterial aerosol dynamics appeared to be very different. For low level PM pollution, the bacterial aerosol structures remained to be relatively constant, in contrast bacterial structures experienced rapid changes during the day not only in abundances but also in species, e.g., several species such as *Psychrobacter* could account for more than 90% of the total abundance as shown in Figure 3 (B). Among the detected species, *Psychrobacter* is a psychrophilic bacterium and used in wastewater treatment(Huang et al., 2018), while *Massilia* is also used in degrading water-borne PAH(Chadhain et al., 2006). The results obtained in this repeated experiment on a different haze day further strengthen the fact that rapid and time-resolved massive changes in bacterial aerosol dynamics have taken place for different PM_2.5_ pollution levels.

Additionally, it is highly interesting that as shown in Figure 4 (A) for the same time, i. e., at 20:00PM on Feb 6, 2018 elevated levels of soluble Na, Ca,Mg, K, Al, Fe and P were observed, which co-occurred with significant bacterial aerosol species shift *(Acinetobacter* increased significantly) as observed in Figure 3 (A). The Ca ion level was observed more than 0.55 μg /m^3^, followed by Na, Mg, K and others as shown in Figure 4 (A). These element ions are necessary for bacterial growth. Likewise, for March 10-13 experiments, several different ion peaks were observed as shown in Figure 4 (B). For those samples collected at 20:00PM of March 11 (“11-20” in the figure) and 08:00AM of March 12 (“12-8” in the figure), elevated Ca, Na and Mg levels were also observed up to 0.9 μg /m^3^ for Ca ion, corresponding to significant bacterial aerosol shifts as observed in Figure 3 (B). It seems that as observed in Figure 3 (B) and Figure 4 (B) *Acinetobacter* and *Massilia* alternated to dominate the bacterial aerosol community structures. When Ca ion level increased *Acinetobacter* started to dominate, while Na level increased *Massilia* started to dominate. The co-occurrence of time-resolved element ion elevation with time-resolved bacterial aerosol structure shift suggest the ions must have been to some extent related to bacterial aerosol dynamics in the air. In addition, we have also constructed a HYSPLIT Trajectory Model for air mass transport for March 10-13, 2018 experiment as shown in Figure 5. As seen from the figure, during the episode air mass was transported from four different directions with different contributions up to 30% of the total transport. In addition, as shown in Figure 5 along the transport directions most of PM_2.5_ levels were below 100 μg/m^3^. Accordingly, it is hard to believe that regional air mass transport contributed directly to the rapid shifts in bacterial aerosol structure shifts, viability and levels.

**Figure 4.**
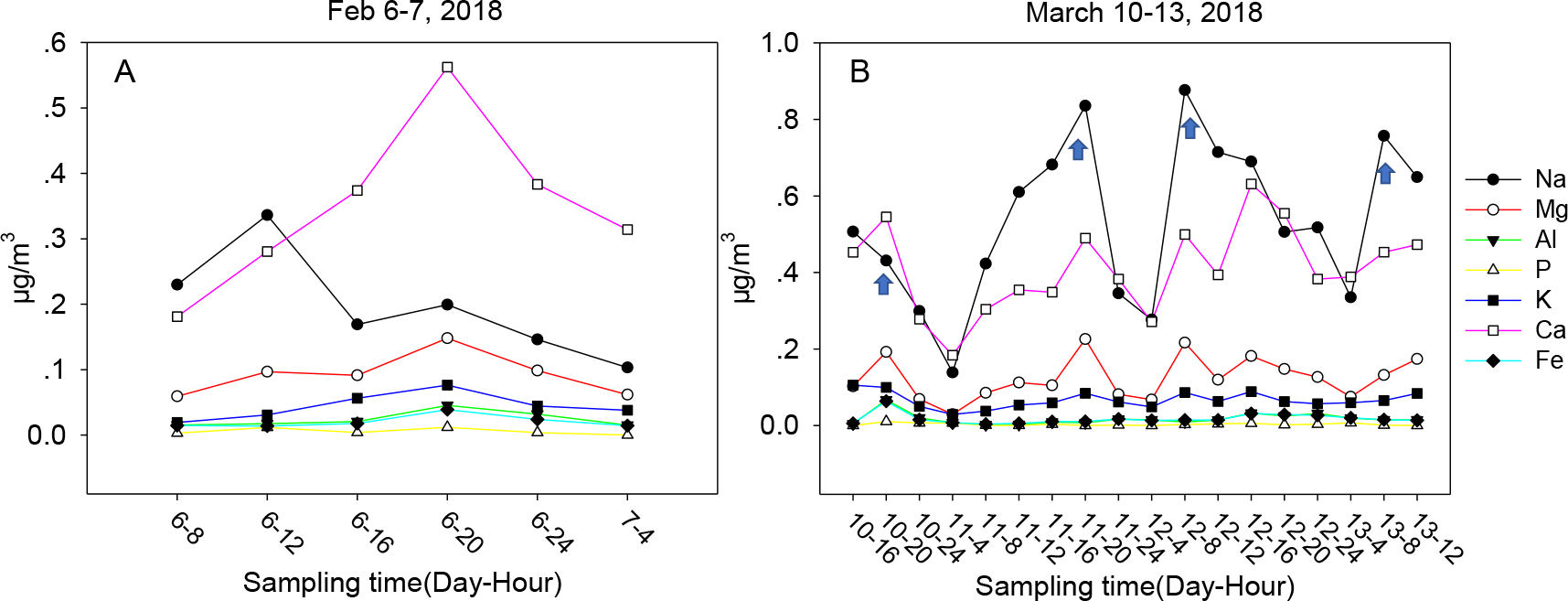
Time-resolved soluble elements (Na, Mg, Al, P, K, Ca, and Fe) concentrations in PM samples collected using the HighBioTrap sampler as described in Figure 3 during Feb 6-7, 2018 and March 10-13, 2018

**Figure 5.**
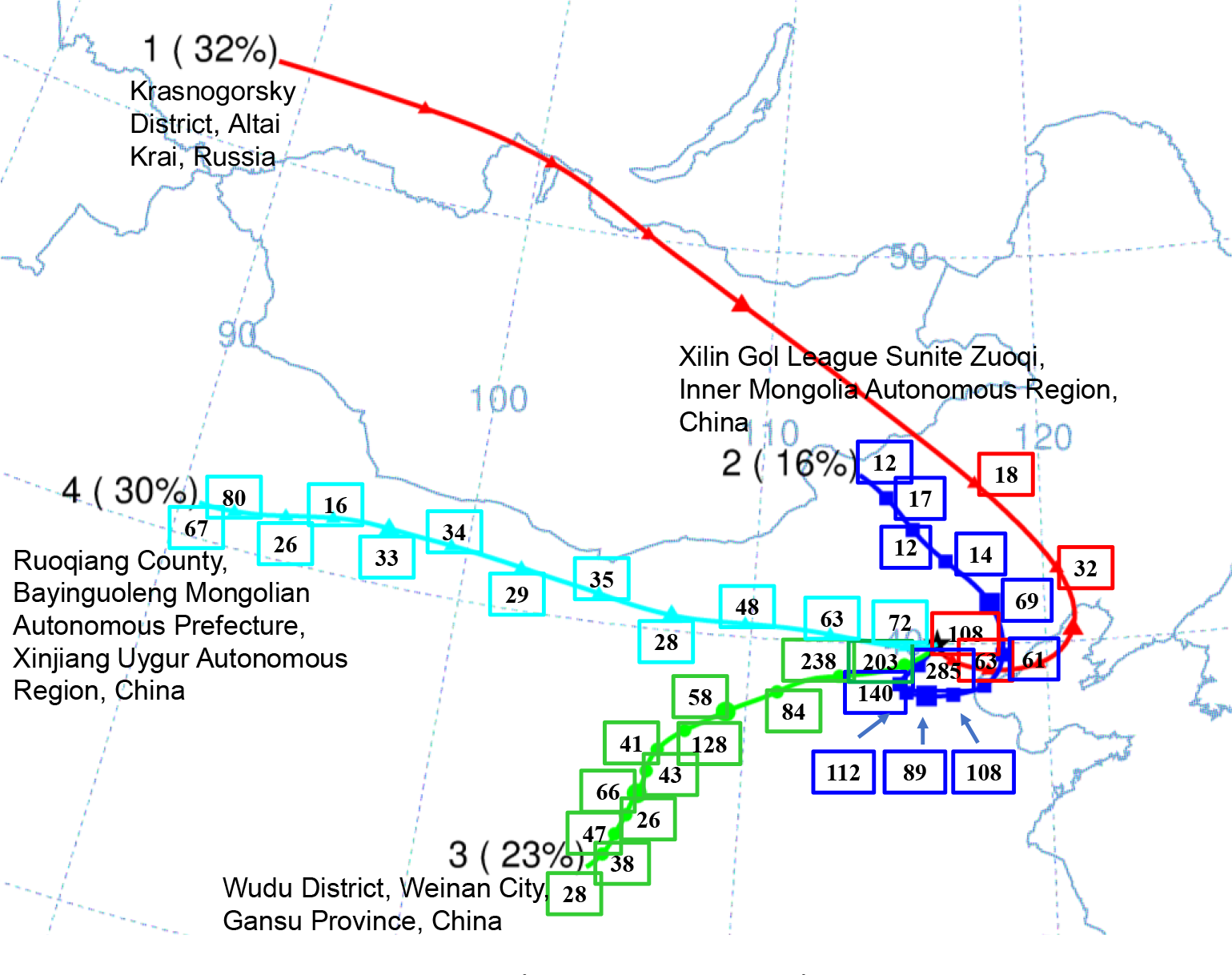
Air mass transport to Beijing (39.99 N 116.32 E) during the March 10-13, 2018 air pollution episodes using HYSPLIT Trajectory Model (NOAA) via Hysplit 4 software. Numbers on four different lines represent different PM_2.5_ mass concentration levels (http://pm25.in/) (μg/m^3^) when the air mass across. The air mass was transported from 4 different directions (1-4) as marked in the figure, accounting for 32%, 16%, 23% and 30% of the total transport contribution (not estimated for its absolute mass amount).

### 4.4 A new frame for understanding air pollution episodes

Air pollution has resulted in tremendous loss of life and economy every year worldwide. Huge amount of research has already been devoted to studying the haze formation and the aerosol chemistry mechanisms(Wang et al., 2006;Zhao et al., 2013;Liu et al., 2013). However, microbial aerosols rarely been considered to play a role in aerosol chemistry especially in highly polluted air. Previously, we have shown using a laser-induced fluorescence-based method that the viable bacterial aerosol concentration was significantly elevated during high PM_2.5_ pollution episodes in Beijing(Wei et al., 2016). Here, for the first time we have captured the size-and time-resolved massive microbial level increase and structure shift events in high PM_2.5_ polluted air in Beijing using a high-volume portable aerosol sampler up to 1000 L/min. Traditionally, people often use low flow rate sampler such as BioSampler (SKC, INc., flow rate=12.5 L/min) and the 4-channel PM sampler (16.7 L/min) or sometimes up to 100 L/min, but most for short time samplings(Gao et al., 2015b;Yuan et al., 2017). These samplers with low sampling rates often fail to capture rapid but transient microbial events in the air since newly bacterial aerosol dynamics could be quickly diluted by open ambient air, and such time-resolved events could also discontinue as atmospheric conditions change from the air. For example, through NMDS analysis we detected a difference in bacterial community structures in the samples collected by the HighBioTrap sampler, but not those collected using the 4-Channel PM sampler (16.7 L/min) as shown in Figure S9 (Supporting Information). Here, by applying a high volume sampler we were able to successfully capture the time-resolved changes in microbial aerosol dynamics in the air.

Accordingly, based on the results obtained here we hypothesize a new haze formation mechanism, i.e., some new aerosol chemistry for highly polluted air as illustrated in Figure 6. As depicted from the figure, bacteria are presumed to be emitted from ground due to various human activities into the air. As such an evidence shown in Figure S10, we found that the bacterial aerosol concentration level exceeded 3000 CFU/m^3^ in a pharmaceutical plant in China. In addition, sequence data shown in Figure S11 (Supporting Information) showed that *Acinetobacter, Psychrobacter, Exiguobacterium*, and *Bacillus* genera were emitted and detected to dominate airborne bacterial community in workshop air inside the pharmaceutical plant. As discussed, these bacteria were also detected in the ambient air in Beijing during the haze episodes, and some of these bacteria experienced rapid concentration changes in the air as presented in Figure 3. Besides, the total VOC concentration for one chimney (pharmaceutical waste treatment workshop) inside the plant was observed to reach a level of 15,030 ppbv, which mainly consisted Acetone (61%) and Acetaldehyde (11%) as seen in Figure S12(Supporting Information). These data support our claim that ground human activities such as pharmacy producing could emit large amounts of bacteria into the air, which were also detected in highly polluted air. As detected in the air samples in this work, the nutrients for bacterial growth such as NH_4_+, NO_3_^−^, Ca, Na, K, Mg, etc. along with other favorable conditions such as RH and gases are readily available in highly polluted atmosphere such as haze episodes in Beijing, accordingly providing required conditions for bacterial growth. Accordingly, we hypothesize that the bacterial growth in the air could have taken place during air pollution episodes as depicted in Figure 6. The bacterial growth typically can undergo for different cycles (in general ∼20-30 min per cycle), depending on the availability of various conditions such as nutrients, RH and temperature. As hypothesized in Figure 6, for typical high PM pollution episodes, atmospheric boundary layer is lowered(Quan et al., 2014), thus leading to increases in ambient pollutant concentrations such as VOC, NH3, NOx, etc. Under these conditions, the airborne bacteria such as *Acinetobacter lwoffii, Bacillus* cereus, *Bacillus pumilus, Exiguobacterium aurantiacum, Arthrobacter ilicis, Bacillus cereus, Bacillus megaterium, Coriobacterium* emitted from ground human activities, e.g., as demonstrated from a pharmaceutical or a wastewater treatment plant, as hypothesized in Figure 6 can play an important role for atmospheric carbon and nitrogen cycles during their growth, thus influencing aerosol chemistry.

**Figure 6.**
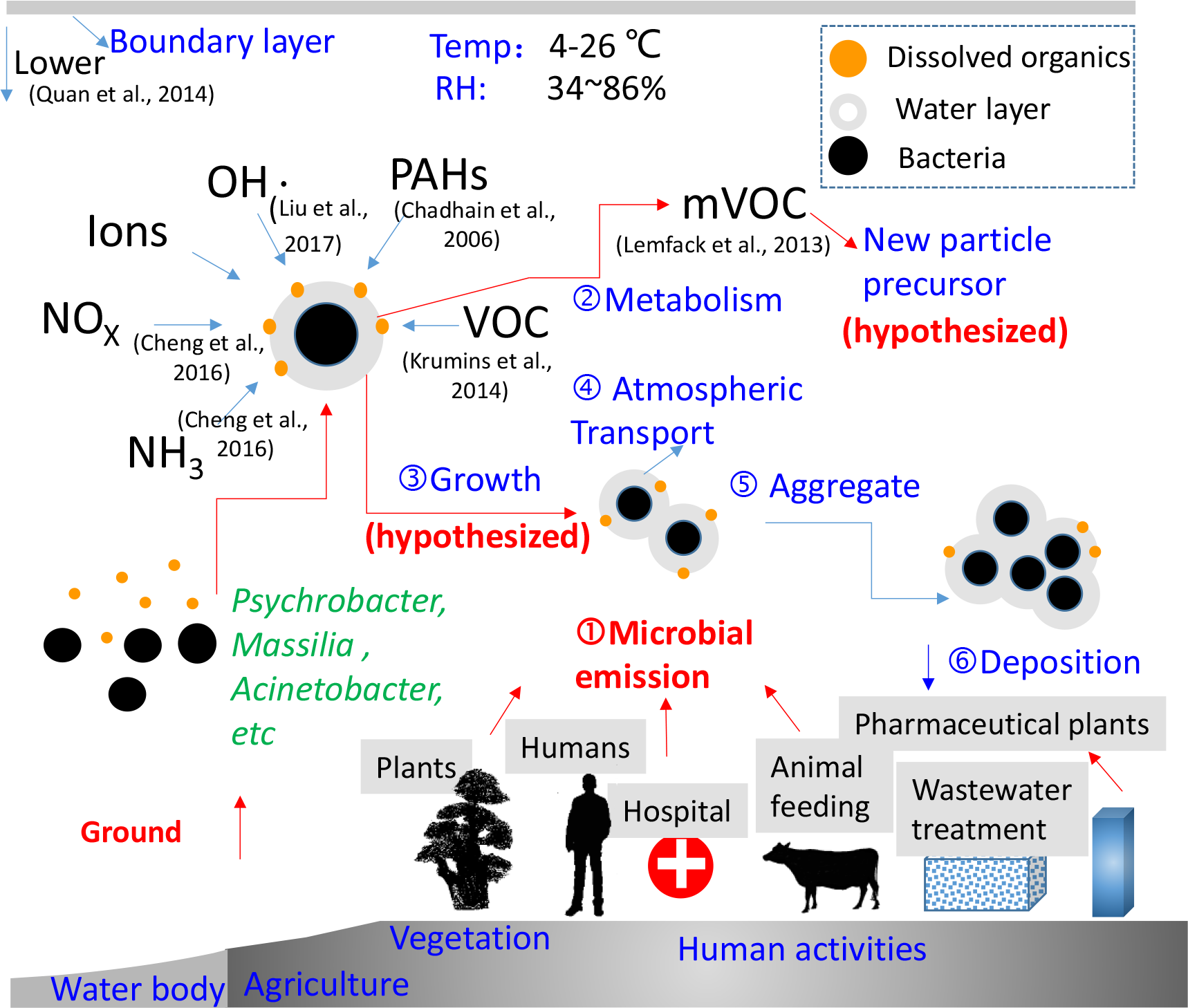
Hypothesized new mechanisms of bacterial growth in the air and possible roles in aerosol chemistry: Ground-emitted bacteria can uptake water vapor, PAH, VOC, NH3, NOx and release mVOCs during the growth; mVOCs could serve as new particle precursors; ultrafine particles (dissolved organics) can attach to the bacterial surface and further mix with water; newly produced bacteria can form aggregates and precipitate.

Studies show that bacteria can use VOC as nutrient(Paavolainen et al., 1998;Krumins et al., 2014), degrade PAH(Chadhain et al., 2006), oxidize NOx(Lambeth, 2004) and scavenge OH radicals(Samake et al., 2017), and at the same time bacteria can also release many mVOCs during their growth(Schulz and Dickschat, 2007). According to the literature, these mVOCs are often small molecular weight molecules (<300 Da) such as hydrocarbons, alcohols, aldehydes and ketones), terpenoids, aromatic compounds, nitrogen containing compounds and volatile sulphur compounds(Lemfack et al., 2013). For example, *Acinetobacter calcoaceticus* can emit 2-oxoethanesulfonic Acid (LogP=−1.18; C_2_H_4_O_4_S) (Schulz and Dickschat, 2007) which is classified as a sulfate and belongs to semi-volatile VOC (sVOC). And *Pseudomonas fluorescens* can emit low volatility Methylsulfonylsulfanylmethane (C_2_H_6_O_2_S_2_, LogP= - 0.06)(Hunziker et al., 2014). In this work, we also detected higher abundance of *Acinetobacter* during higher PM pollution episodes. On the other hand, *Ralstonia solanacearum* can emit nitrogen containing compounds such as 2-chloro-2-nitropropane (C_3_H_6_ClNO_2_)(Spraker et al., 2014). In our work, we have detected higher abundance of *Ralstonia* species in our work as shown in Figure 3 A). In another work, it was shown that *Psychrobacter* spp. can emit 2,3-Dimethyl-oxirane, 2-Butanone, 2-Formylhistamine, 2-Methyl-2-propanol, Acetaldehyde, Acetone, Ethylene oxide, Isopropylalcohol, and Trimethylamine(Broekaert et al., 2013). For a complete reference to mVOC emitted by both bacteria and fungi, there is a database available in the literature(Lemfack et al., 2013). As seen from the database, bacteria, depending on the species type, could emit mVOCs containing sulfur, nitrogen or simply hydrocarbon^35^. These smaller molecules (mVOCs) can be further oxidized into ultrafine particles under oxidizing environments, and some of them are sVOC with lower vapor pressure.

These analyses led to our belief that the bacterial growth, accordingly emitting mVOCs as new particle precursors, in the air could play a very important role for new particle formation as hypothesized in Figure 6. From the literature, we have found that new particle formation occurred during the early mornings (04:00-06:00) or during the afternoon time period (18:00-22:00) (Liu et al., 2008;Wang et al., 2014). During these same time periods, massive changes in bacterial aerosol levels and structure shifts were also observed in our work as shown in Figure 1-3. As illustrated in Figure 6, these ultrafine particles eventually could grow into fine particles via hydroscopic growth or through atmospheric oxidization(Guo et al., 2014). Depending on the bacterial species or the ambient pollutants, major components of the sVOC ultrafine particles emitted if any in the air as hypothesized could be organic compounds, nitrogen containing or sulfates, thus likely impacting haze episode dynamics. Simultaneously, airborne bacteria can also scavenge OH radicals(Samake et al., 2017) and react with metals such as Fe(Chen et al., 2013) in the water layer as depicted in Figure 6. During a certain time period, the available nutrients for bacteria are depleting in the atmosphere, then the bacterial growth may slow down and eventually stop. From our observations, these bacteria are mostly in the size range 0.5-1 μm, and those newly produced ones can be quickly attached by atmospheric ultrafine particles or form aggregates, which could later grow into bigger particles and settle to the ground. When the latter occurs, the high PM pollution episode is disappearing. Here, some psychrophilic bacteria were also detected, indicating microbial growth in the cold air could also occur, e.g., during the winter periods when frequent hazes occur. Those bacteria trained for engineering purposes can also significantly impact human health and ecology when they have undergone growth if any in the air and further precipitated into the local environments or inhaled by humans during high air pollution episodes. For example, we have detected abundant human pathogen such as *Acinetobacter lwoffii* in highly polluted air, which thus presents a significant public health concern. The hypothesized bacterial growth in the air partly could explain the underestimates (e.g. HCHO was underestimated by about 16% compared to observation) of atmospheric VOCs by current models, and also the under-prediction (about 30%) of OH radical loss rate (OH radicals react with VOCs)(Whalley et al., 2016). Here, we only investigated the bacterial dynamics, while those airborne fungal species could also have an impact in the air under favorable conditions. This work was mainly conducted in Beijing, and certainly many other urban settings are warranted to further verify the findings from this work. The results from this work provide a new understanding of current aerosol chemistry known so far especially in highly polluted air. The relevant results might find their values in future cost-effective air pollution control practices. The hypothesized airborne bacterial growth mechanism can be explored using an isotope-based experimental method in future studies.

## 5. Acknowledgements

This study was supported by the NSFC Distinguished Young Scholars Fund Awarded to M. Yao (21725701), and the National Natural Science Foundation of China (Grants 91543126, 21611130103, 21477003, 41121004), the Ministry of Science and Technology (grants 2016YFC0207102, 2015CB553401, and 2015DFG92040).

## 6. Author contribution

MY designed the study, TZ performed the experiments, XL, MW helped conducting some of the samplings, XL, HC collected the air samples from the pharmaceutical plant. TZ and MY prepared the manuscript with contributions from all co-authors. The results here only represent scientific evidences and the authors’ understanding of the problem.

