## Supplemental file for "Time- and size-resolved bacterial aerosol dynamics in highly polluted air: new clues for haze formation mechanism"

Submitted to

***BioRxiv***

* Corresponding Author:
Maosheng Yao, PhD
Boya Professor
State Key Joint Laboratory of Environmental Simulation and Pollution Control,
College of Environmental Sciences and Engineering,
Peking University, Beijing 100871, China

Ph: +86 01062767282

Jan 6, 2018

Beijing, China

### Table of Contents

**Supplementary Text** Experimental protocols of sample preparation and sequencing

**Supplementary Figures**

**Figure S1** Soluble element concentrations in the size range of 0.56-1 µm and 0.32 µm -0.56µm of the NanoMoudi samples during Sept 11 to 14, 2017.

**Figure S2** The $O_3$, $NO_2$, $SO_2$, CO concentration levels measured in Beijing during Sept 11-14,2017. These pollutant data were obtained from https://www.aqistudy.cn/ (accessed on July 7, 2018)

**Figure S3** $K^+_{BB}$ concentrations in the size range of 0.56-1 µm and 0.32-0.56 µm.

**Figure S4** Bacterial DNA stain (BackLight method as described in the experimental section) results from air samples in the size range of 0.56-1 µm and 0.32-0.56 µm on Sept 11 to 14, 2017.

**Figure S5** Viable and dead bacteria percentages in different size ranges of 10 nm-20 µm from the NanoMoudi samples collected during Sept 11 to 14, 2017.

**Figure S6** Total bacteria percentages in different size range in the NanoMoudi samples collected for both Days and Nights during Sept 11-14,2017.

**Figure S7** Day and night endotoxin concentrations in size-fractionated particulate matters, collected during Sept 11 to 14, 2017.

**Figure S8** $PM_{2.5}$ concentration at different times on May 12, 16:00 – May 13, 22:00. Information was obtained from https://www.aqistudy.cn/ (accessed on July, 2018).

**Figure S9** NMDS analysis for samples collected on March 10-12, 2018.

**Figure S10** The bacterial and fungal aerosol concentration levels in a pharmaceutical plant and nearby locations in a Chinese city.

**Figure S11** Culturable bacterial community structures of the air samples collected using the HighBioTrap sampler inside a pharmaceutical plant in a Chinese city in 2017.

**Figure S12** Various VOCs were detected in the air samples at the chimneys and workshops of a pharmaceutical plant inside a pharmaceutical factory in a Chinese city.

**Supplementary Tables**

Table S1 Sampling information (sampling dates, duration and the sampler)

**References**

### Supplementary Materials

#### Supplementary Text

From March 26 to 30, 2018, eight air samples were collected for day and night time using a 4-channel particulate matter sampler (TH-16A, Wuhan Tianhong Instruments Co., Ltd., China) on teflon filters at a flow rate of 16.7 L $min^{-1}$ on Peking University campus. Specific sampling times and meteorological conditions are provided in the Table S1.

**PM-borne reactive oxygen species (ROS) and metal analysis using DTT assay and ICP-MS**

The DTT (DL-Dithiothreitol) assay procedure in this study was performed according to the process described in a previous work[1], except that we added 100 μL samples solution and double distilled water in the reaction mixture. A total of the 12 sample extracts from particulate matter collected onto stage 6 and 7 of the NanoMoudi and the negative control sample were filtrated through by 0.22 μm pore-size filters for the analysis of 24 metal elements (Na, P, K, Ca, Ti, V, Cr, Co, Ni, As, Mo, Cd, Tl, Mg, Al, Fe, Zn, Mn, Cu, Se, Ba, Pb, Th, and U) using ICP-MS (aurora M90, Analytikjena, German). The blank membrane was also used for analysis as a negative control.

**The PM-borne total live/dead bacteria analysis using DNA stain method**

To study the viability of bacteria in the particulate matters, a live/dead viability kit

(L7012 BacLight Viability Kit, Invitrogen) was used in this study. Two dyes were included in this kit: SYTO 9 labels bacteria with intact membranes green, and propidium iodide labels dead or injured bacteria red. For each of 78 samples, 300 µl (NanoMoudi Samples) or 500 µl (HighBiotrap samples) extraction solution was used together with 1 negative control sample and 1 positive control sample (*Bacillus subtilis*), respectively, for DNA stain. Then, we added pre-mixed fluorescent dyes (SYTO® 9 and propidium iodine, 1:1) to the samples with a 3:1000 (V: V) ratio. After 15min incubation in the dark, each mixture was filtered and concentrated on a black hydrophilic polycarbonate membrane filter (Millipore, GTBP02500, 25 mm, pore size=0.2 µm). Each mixture would form a circle of about 80 mm in diameter on the filters. A fluorescence microscope (BX63 with DP27-CU camera, Olympus, Japan) was applied to view and photograph the DNA stained samples. For each sample, at least five pictures were taken for randomly selected microscopic view fields (field area: 135 µm × 75 µm). The number of red or green cells were manually counted for each microscopic view photo, and relevant airborne viable or dead bacterial aerosol concentration level was determined by considering the amount of air sampled, volume of sample extract used, and the area of microscopic view area using the following equation.

$$\mathrm{B} = \frac{NA}{\pi r^2 v_1} v_2 / v_a$$

B: Bacteria concentration, cell/m$^3$

N: The average number of red or green cells in each microscopic view photo

A: Microscopic view area, m$^2$

r: Circle radius on the filter, m

$v_1$: Sample extract used for membrane filtration, mL

$v_2$: Whole sample extract volume, mL

$v_a$: Sampling air volume, $m^3$

**Bacterial species identification using VITKE MS and MICROFLEX**

Here, LB agar (tryptone 10 g/L, yeast extract 5 g/L, NaCl 10 g/L, agar 15 g/L) was used for the cultivation of the air samples. For each agar plate, 100 µl air sample extract solution was plated and incubated at 37◦C for 48 h. Two blank plates were also placed in the incubator as negative controls. After the cultivation, the colony forming units (CFU) were individually picked using a sterilized loop. All single colonies were identified both by VITKE MS (compared with SARAMIS VERSION 4.14 bank) and MICROFLEX (compared with BIOTYPER VERSION 4.0 bank). Blank agar plates were also incubated and used to eliminate the possibility agar contamination.

**PM-borne Endotoxin assay using the LAL assay, bacterial viability using DNA stain method, and reactive oxygen species (ROS) using DL-Dithiothreitol (DTT) assay**

The samples collected from the NanoMoudi were first centrifuged at 4℃ at 3000 rpm in 10 min, and the supernatants were then transferred to the new tubes for the Limulus amebocyte lysate (LAL) assay. A 5-point standard curve from 0.005 to 50.0 EU/ml was generated using 10-fold serial dilutions of endotoxin standards (Control Standard Endotoxin, CSE, Associate of Cape Cod, Inc., Eastham, MA, USA) by endotoxin-free water (LAL Reagent Water, Associate of Cape Cod, Inc., Eastham, MA,

USA). The LAL agent (Associate of Cape Cod, Inc., Eastham, MA, USA） diluted in the Glucashield Beta Glucan Inhibiting Buffer (Associate of Cape Cod, Inc., Eastham, MA, USA) was used in the assay according to the manufacturer's guidelines. The standards, negative controls and air samples were pre-incubated and analyzed using a microplate reader running SoftMaxPro 5.4.1 software (SpectraMax 340; Molecular Devices, Sunnyvale, CA) with photometric measurements taken at 37°C every 1min for 60 min at 405 nm. The viability of bacteria in the particulate matters was studied using a live/dead viability kit (L7012 BacLight Viability Kit, Invitrogen) as described in Supplementary Text. PM-borne reactive oxygen species (ROS) and metal analysis were conducted using DTT assay and ICP-MS as documented in Supplementary Text.

**Total PM-borne bacteria and community structure analysis using molecular methods**

For detecting the total bacteria, 78 samples and 2 negative control samples (blank membrane samples) were extracted for DNA using the bacteria DNA Extraction Kit (Tiangen Biotech, Inc., Beijing, China) according to the manufacturer's guidelines. Extracted 200 μl DNA samples were stored at -20℃ until qPCR analysis. The concentrations of 16S rDNA in the particles of 13 different sizes from the NanoMoudi were quantified using the quantitative PCR (qPCR) in this study. In a total 50 μl reaction volume for the qPCR experiments, the primers and probe used for universal qPCR assays were: forward primer: 5'-TCCTACGGGAGGCAGCAGT-3' (Tm=59±4 °C), reverse primer: 5'-GGACTACCAGGGTATCTAATCCTGTT-3' (Tm=58±1 °C), and probe: (6-FAM)-5'-CGTATTACCGCGGCTGCTGGCAC-3'-(TAMRA) (Tm=69±9 °C)[2]. The cycle conditions were: 94 °C for 3 min, followed by 5 cycles at 94 °C for 30 s, 45 °C for 30 s, and 62 °C for 30 s, and 20 cycles at 94 °C for 20 s, 55 °C for 20 s, and 72 °C for 30 s, and a final extension at 72 °C for 5 min [2]. In addition, the air samples collected both by the NanoMoudi and the HighBioTrap were also sequenced for V3–V4 region of bacteria 16S ribosomal RNA genes for analyzing the bacterial aerosol community structures by Illumina platforms (Sangon Biotech, Inc., Shanghai, China) according to a previously described protocol[3].

### Supporting Figures

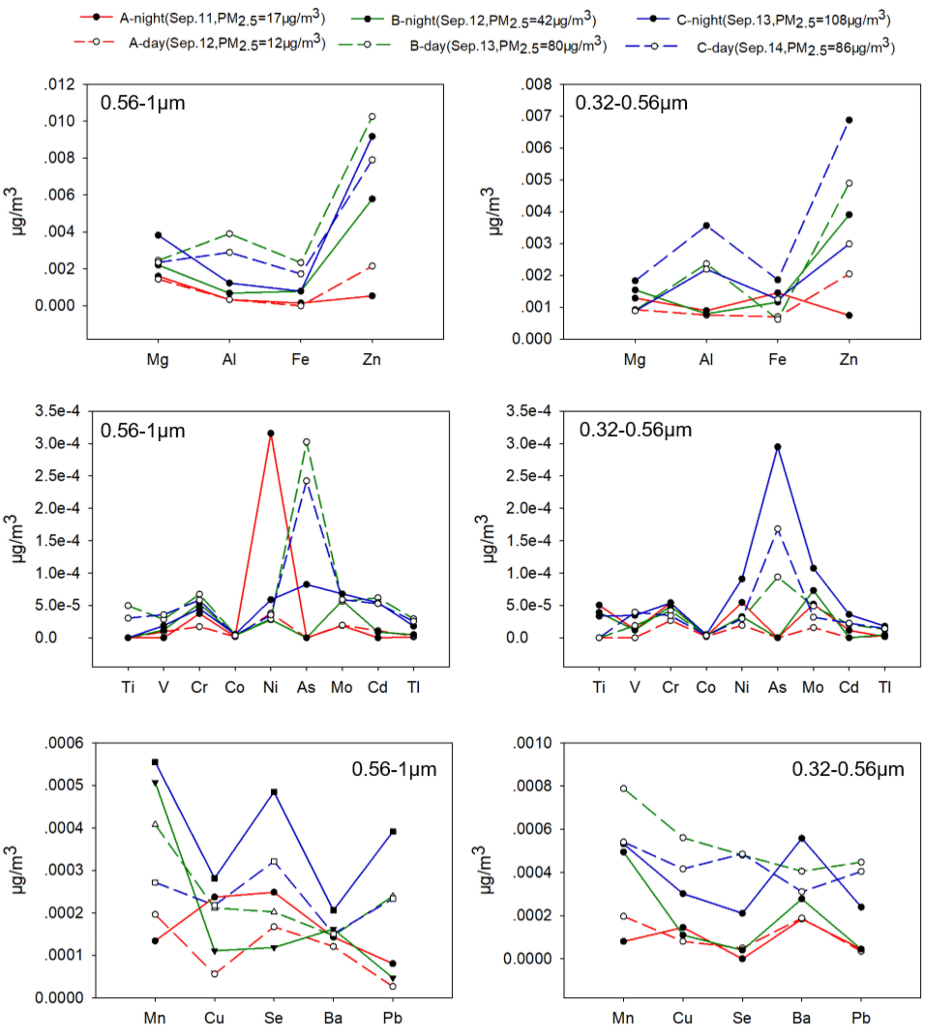


**Figure S1** Soluble element (Ti, V, Cr, Co, Ni, As, Mo, Cd, Tl, Mg, Al, Fe, Zn, Mn, Cu, Se, Ba, Pb) concentrations in the size range of 0.56 -1μm (the sixth stage) and 0.32 -0.56μm (the seventh stage) of the NanoMoudi samples, as described in Figure 1, during Sept 11 to 14, 2017.

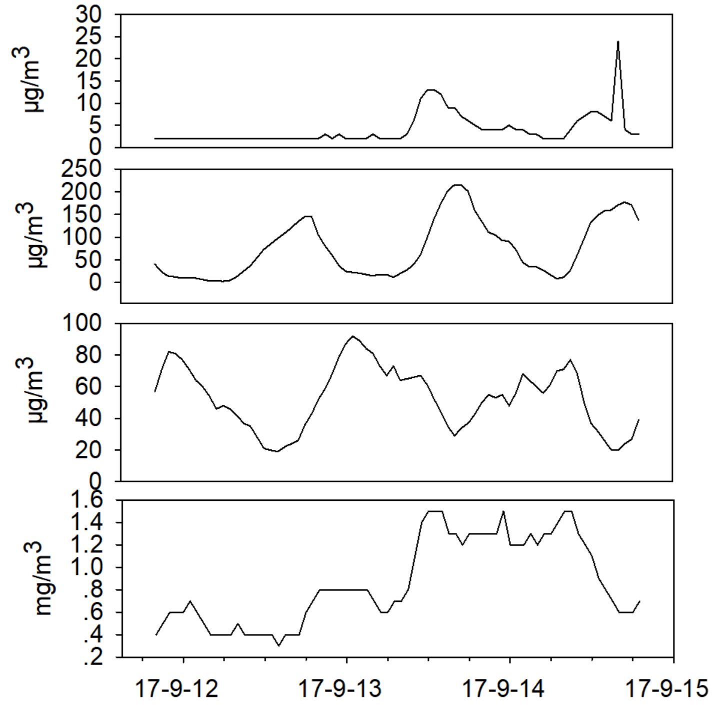


**Figure S2** The $O_3$, $NO_2$, $SO_2$, CO concentration levels measured in Beijing during Sept 11-14,2017. These pollutant data were obtained from https://www.aqistudy.cn/ (accessed on July 7, 2018)

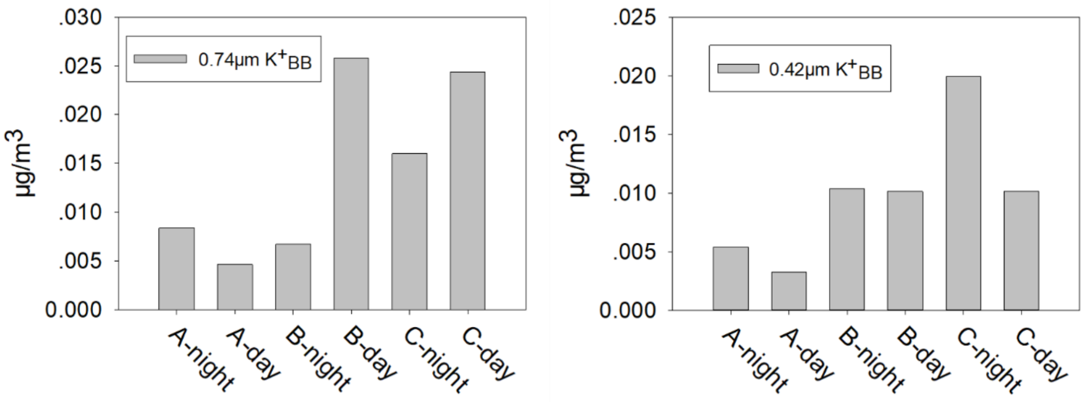


**Figure S3** $K^+_{BB}$ concentrations in the size range of 0.56-1μm (0.74 μm in the figure) and 0.32-0.56μm (0.42 μm in the figure), as described in Figure 1 and Figure S1, on Sept 11 to 14, 2017. The previously reported equation for K contribution from the biomass burning,1 $K^+_{BB}=(K^+-0.036*Na^+-0.12*Ca^{2+})/0.988$, is used here[4]. A, B and C represents three different days： Sept 11, Sept 12, and Sept 13, 2017, respectively. Night refers to 20:00PM-6:00AM, and Day refers to 7:00AM-19:00PM in the figure.

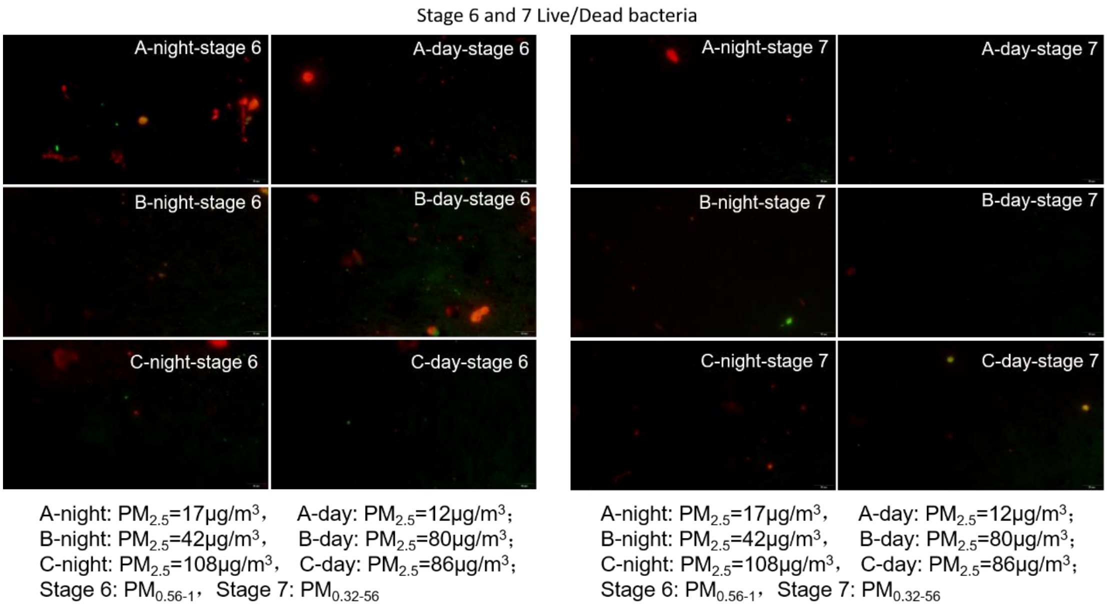


**Figure S4** Bacterial DNA stain (BackLight method as described in the experimental section) results from air samples in the size range of 0.56-1 μm and 0.32-0.56 μm, as described in Figure 1 and Figure S1, on Sept 11 to 14, 2017. Stage 6 refers to the NanoMoudi cutoff stage of 0.56 -1μm and Stage 7 refers to the cutoff stage of 0.32 - 0.56 μm. A, B and C represents three different days：Sept 11, Sept 12, and Sept 13, 2017, respectively. Night refers to 20:00PM-6:00AM, and Day refers to 7:00AM-19:00PM in the figure.

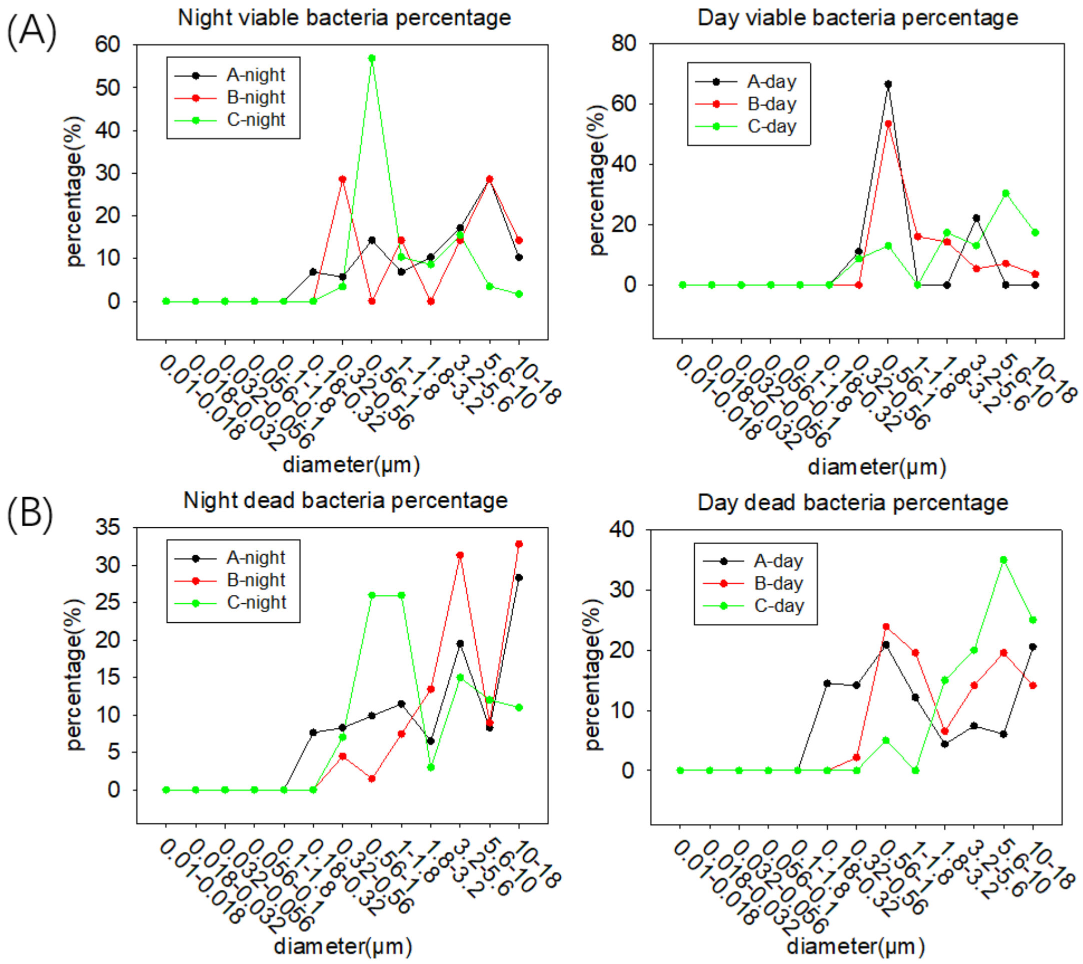


**Figure S5** Viable (A) and dead bacteria (B) percentages in different size ranges of 10nm-20 μm from the NanoMoudi samples, as described in Figures S1 and Figure 1, collected during Sept 11 to 14, 2017. A, B and C represents three different days：Sept 11, Sept 12, and Sept 13, 2017, respectively. Night refers to 20:00PM-6:00AM, and Day refers to 7:00AM-19:00PM in the figure.

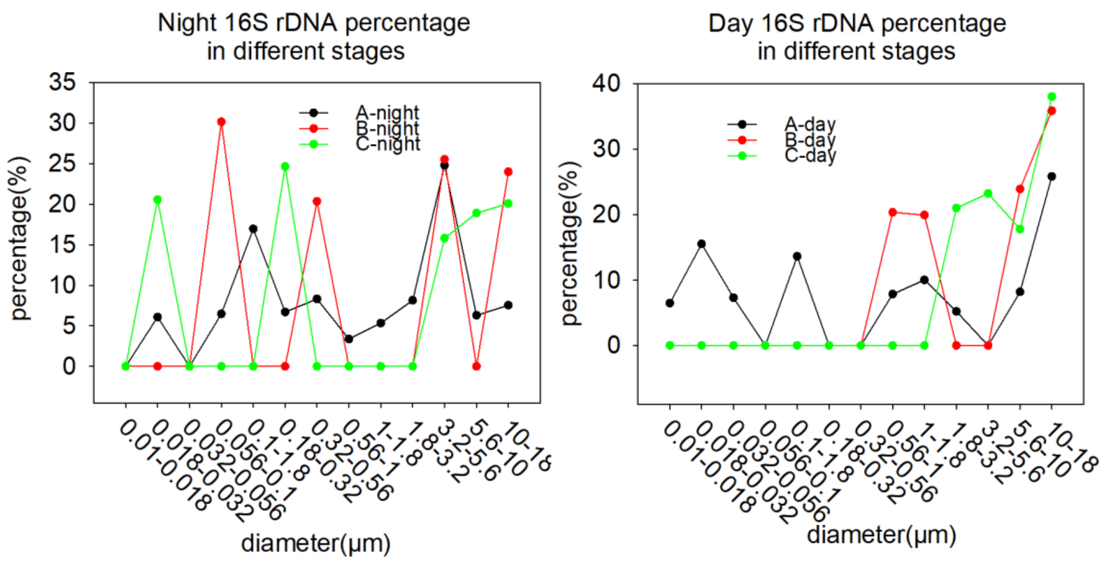


**Figure S6** Total bacteria percentages in different size range in the NanoMoudi samples collected for both Days and Nights during Sept 11-14,2017. The bacterial concentrations were quantified using qPCR as described in the experimental section. A, B and C represents three different days： Sept 11, Sept 12, and Sept 13, 2017, respectively. Night refers to 20:00PM-6:00AM, and Day refers to 7:00AM-19:00PM in the figure.

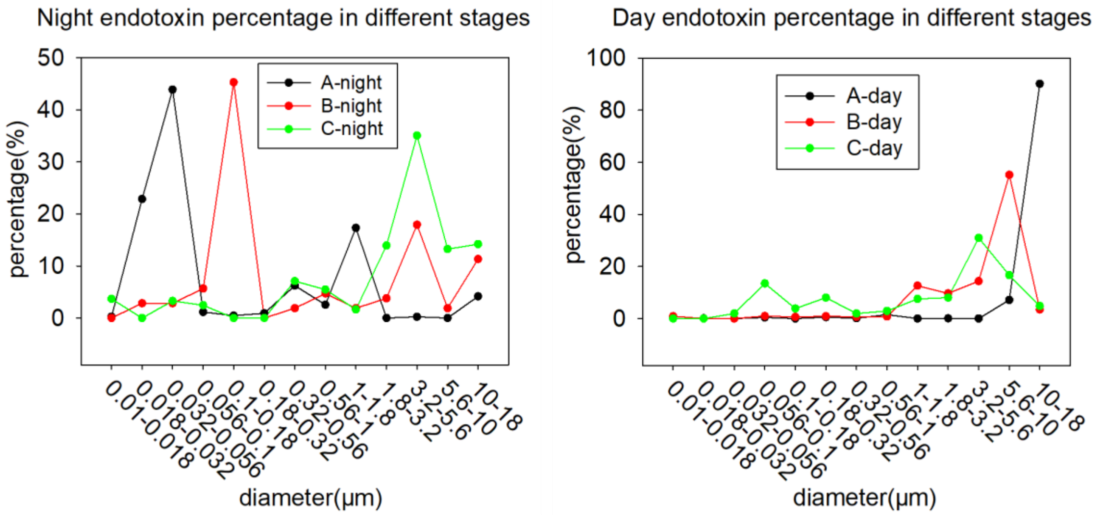


**Figure S7** Day and night endotoxin concentrations in size-fractionated particulate matters, collected during Sept 11 to 14, 2017. A, B and C represents three different days： Sept 11, Sept 12, and Sept 13, 2017, respectively. Night refers to 20:00PM-6:00AM, and Day refers to 7:00AM-19:00PM in the figure. Air samples, as described in Figure 1, were collected into 13 different stages with cutoff sizes of 10nm-20 μm using the NanoMoudi at a flow rate of 28 L/min for 12 hours during the day (7:00 AM to 19:00PM) and 10 hours during the night (20:00PM to 6:00AM) on Sept 11-13, 2017.

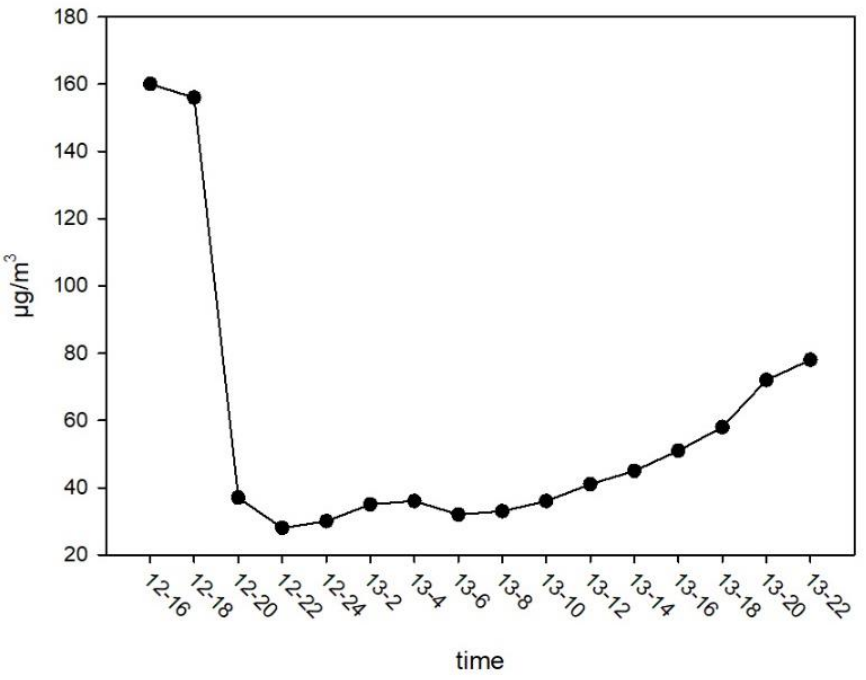


**Figure S8.** $PM_{2.5}$ concentration at different times on May 12, 16:00 – May 13, 22:00.

Information was obtained from https://www.aqistudy.cn/ (accessed on July 7, 2018).

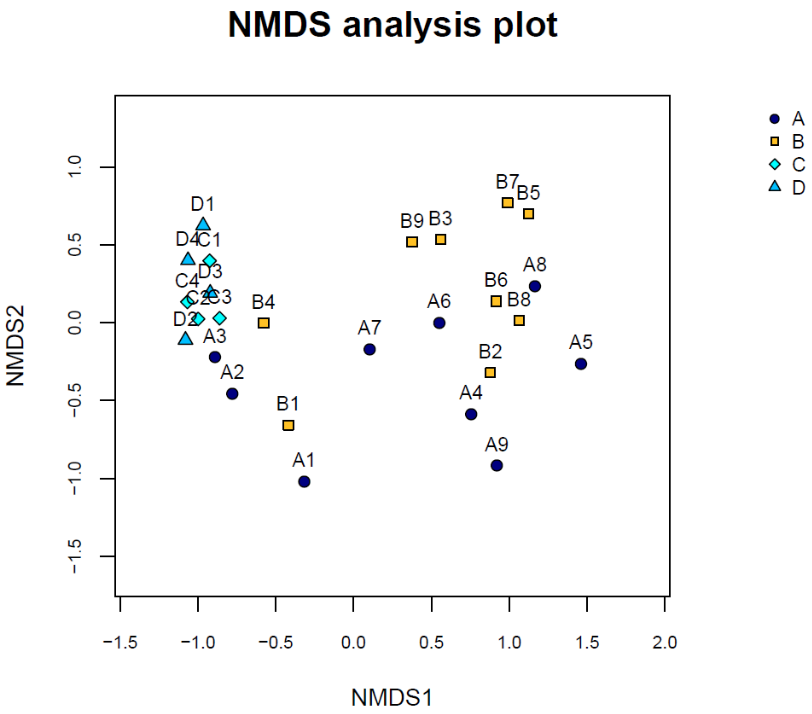


**Figure S9** NMDS analysis for samples collected in March 10-12, 2018. In the figure, A represents samples collected using the high flow sampler HighBioTrap at day-time, March 10-12, 2018, 20 min sampling each, every 4 hours; B represents samples collected by the HighBiotrap (1000 L/min) at night-time on March 10-12, 2018, C represents samples collected using 4-Channel Particulate Matter Sampler (Wuhan Tianhong Instruments Co., Wuhan, China) at day-Time continuously for 12h, March 27-30, 2018; D represents samples collected by 4-Channel Particulate Matter Sampler at night-time continuously for 12h, March 27-30, 2018. Bacterial structures in PM were detected to be different between the samples collected through the 4-Channel Sampler and the HighBiotrap Sampler. In addition, airborne bacterial structures difference in the samples collected by the HighBioTrap was detected between the day and night, but not for the samples collected using the 4-Channel Sampler based on the NMDS analysis.

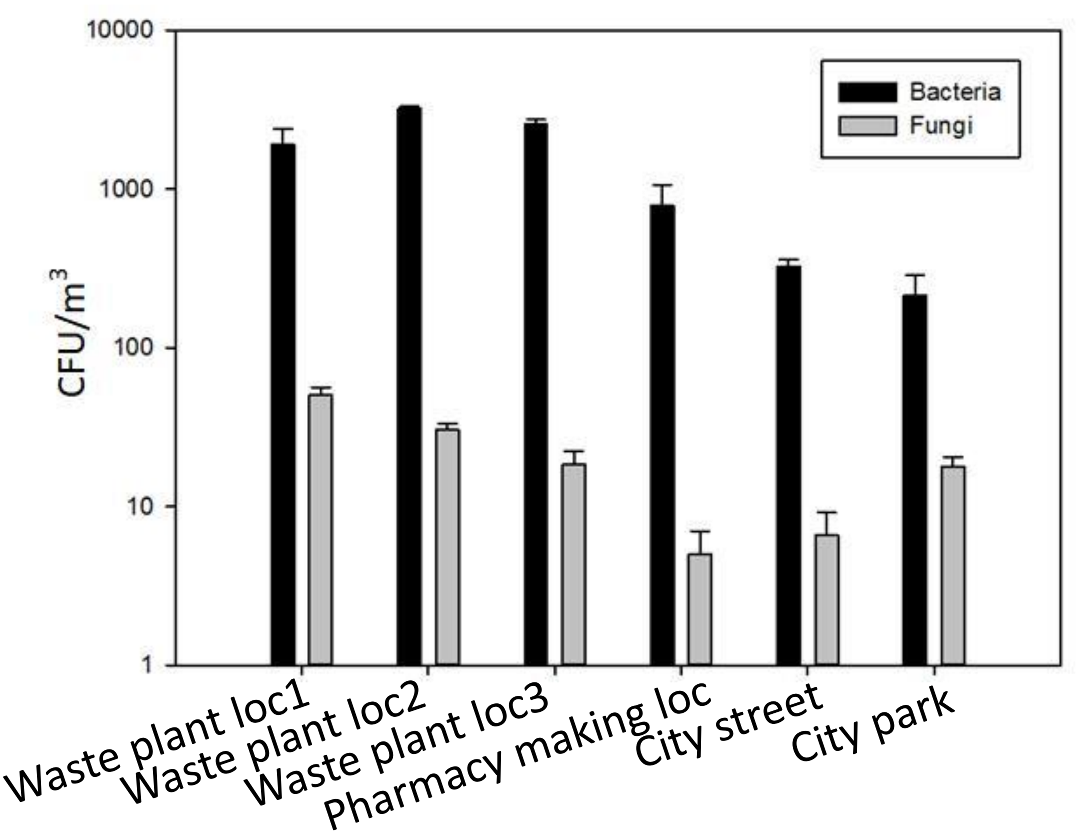


**Figure S10** The bacterial and fungal aerosol concentration levels in a pharmaceutical plant and nearby locations in a Chinese city. Three samples for each location were collected using the HighBioTrap sampler at an air flow rate of 1000 L/min for 20 min at each location in March, 2017. Data points represent the averages and standard deviations from three repeats.

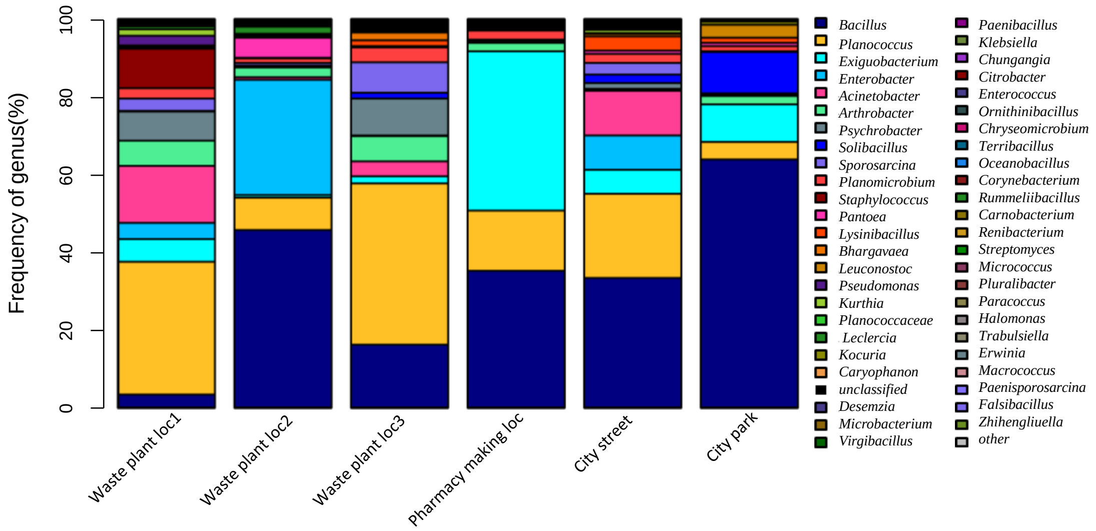


**Figure S11** Culturable bacterial community structures of the air samples collected using the HighBioTrap sampler inside a pharmaceutical plant in a Chinese city. Samples were collected in May, 2017.

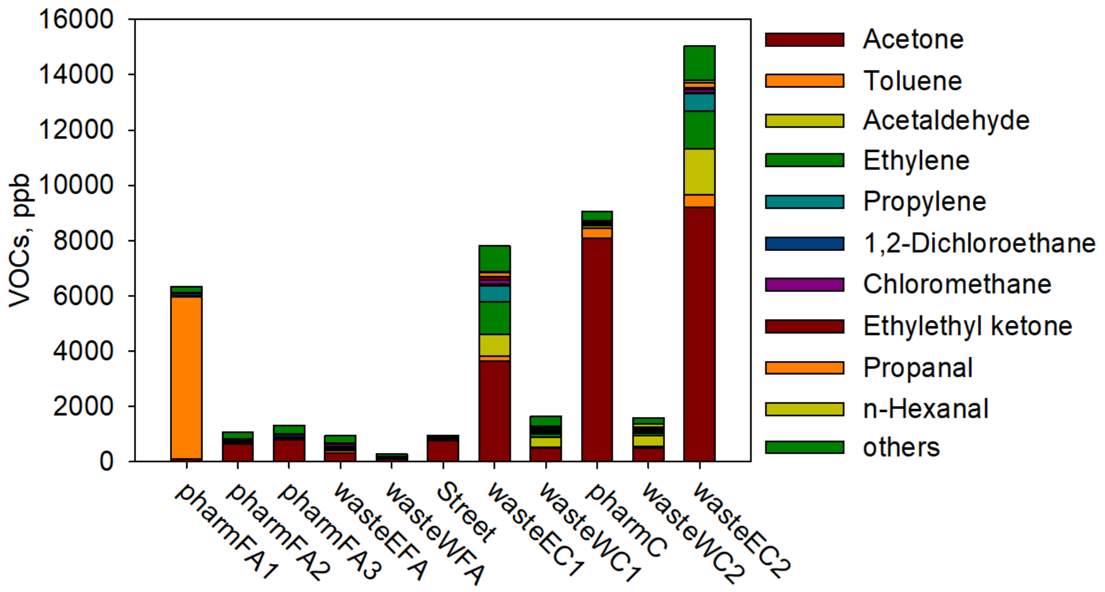


**Figure S12** Various VOCs were detected in the air samples at the chimneys and workshops of a pharmaceutical plant inside a pharmaceutical factory in a Chinese city. Among the VOC sampling sites, pharmFA refers to the pharmacy workshops (indoor rooms), wasteFA refers to the pharmacy waste treatment workshops (indoor rooms), and wasteWC refers to the chimney of pharmaceutical waste treatment workshop. VOC air samples were collected using Silonite™ Classical Canisters (Entech Instruments, Simi Valley, CA 93065) in May 2017, and further analyzed by a commercial company (Wuhan Tianhong Instruments Co., Ltd.)

**Table S1 Sampling information (sampling dates, duration and the sampler)**

| Sampling site | Start time | End time | Sampling time | Sampler |
|---|---|---|---|---|
| The main campus of Peking University (39°59’N E116°18’) | 2017.09.11 20:00 | 2017.09.12 06:00 | 12h | NanoMoudi |
| | 2017.09.12 07:00 | 2017.09.12 19:00 | 12h | |
| | 2017.09.12 20:00 | 2017.09.13 06:00 | 12h | |
| | 2017.09.13 07:00 | 2017.09.13 19:00 | 12h | |
| | 2017.09.13 20:00 | 2017.09.14 06:00 | 12h | |
| | 2017.09.14 07:00 | 2017.09.14 19:00 | 12h | |
| | 2018.02.06 08:00 | 2018.02.06 09:00 | 20min X 3 | HighBioTrap |
| | 2018.02.06 12:00 | 2018.02.06 13:00 | 20min X 3 | |
| | 2018.02.06 16:00 | 2018.02.06 17:00 | 20min X 3 | |

| | | |
|---|---|---|
| 2018.02.06 20:00 | 2018.02.06 21:00 | 20min X 3 |
| 2018.02.07 00:00 | 2018.02.07 01:00 | 20min X 3 |
| 2018.02.07 04:00 | 2018.02.07 05:00 | 20min X 3 |
| 2018.03.10 16:00 | 2018.03.10 16:40 | 20min X 2 |
| 2018.03.10 20:00 | 2018.03.10 20:40 | 20min X 2 |
| 2018.03.11 00:00 | 2018.03.11 00:40 | 20min X 2 |
| 2018.03.11 04:00 | 2018.03.11 04:40 | 20min X 2 |
| 2018.03.11 08:00 | 2018.03.11 08:40 | 20min X 2 |
| 2018.03.11 12:00 | 2018.03.11 12:40 | 20min X 2 |
| 2018.03.11 16:00 | 2018.03.11 16:40 | 20min X 2 |
| 2018.03.11 20:00 | 2018.03.11 20:40 | 20min X 2 |

| | | | |
|---|---|---|---|
| 2018.03.12<br>00:00 | 2018.03.12<br>00:40 | 20min X 2 | |
| 2018.03.12<br>04:00 | 2018.03.12<br>04:40 | 20min X 2 | |
| 2018.03.12<br>08:00 | 2018.03.12<br>08:40 | 20min X 2 | |
| 2018.03.12<br>12:00 | 2018.03.12<br>12:40 | 20min X 2 | |
| 2018.03.26<br>19:40 | 2018.03.27<br>07:40 | 12h | |
| 2018.03.27<br>07:45 | 2018.03.27<br>19:47 | 12h | |
| 2018.03.27<br>19:55 | 2018.03.28<br>07:55 | 12h | |
| 2018.03.28<br>08:00 | 2018.03.28<br>20:00 | 12h | TH-16A |
| 2018.03.28<br>20:06 | 2018.03.29<br>07:36 | 12h | |
| 2018.03.29<br>07:40 | 2018.03.29<br>19:11 | 12h | |
| 2018.03.29<br>19:41 | 2018.03.30<br>07:11 | 12h | |

| | | | |
|---|---|---|---|
| 2018.03.30 07:41 | 2018.03.30 19:11 | 12h | |
| 2018.05.12 16:00 | 2018.05.12 16:40 | 20min X 2 | |
| 2018.05.12 18:00 | 2018.05.12 18:40 | 20min X 2 | |
| 2018.05.12 20:00 | 2018.05.12 20:40 | 20min X 2 | |
| 2018.05.12 22:00 | 2018.05.12 22:40 | 20min X 2 | |
| 2018.05.13 00:00 | 2018.05.13 00:40 | 20min X 2 | HighBioTrap |
| 2018.05.13 02:00 | 2018.05.13 02:40 | 20min X 2 | |
| 2018.05.13 04:00 | 2018.05.13 04:40 | 20min X 2 | |
| 2018.05.13 06:00 | 2018.05.13 06:40 | 20min X 2 | |
| 2018.05.13 08:00 | 2018.05.13 08:40 | 20min X 2 | |
| 2018.05.13 10:00 | 2018.05.13 10:40 | 20min X 2 | |

| | | |
|---|---|---|
| 2018.05.13 12:00 | 2018.05.13 12:40 | 20min X 2 |
| 2018.05.13 14:00 | 2018.05.13 14:40 | 20min X 2 |
| 2018.05.13 16:00 | 2018.05.13 16:40 | 20min X 2 |
| 2018.05.13 18:00 | 2018.05.13 18:40 | 20min X 2 |
| 2018.05.13 20:00 | 2018.05.13 20:40 | 20min X 2 |
| 2018.05.13 22:00 | 2018.05.13 22:40 | 20min X 2 |
